# The estrous cycle modulates early-life adversity effects on mouse avoidance behavior through progesterone signaling

**DOI:** 10.1101/2022.02.23.481634

**Authors:** Blake J. Laham, Sahana S. Murthy, Monica Hanani, Mona Clappier, Sydney Boyer, Betsy Vasquez, Elizabeth Gould

## Abstract

Early-life adversity (ELA) predisposes individuals to develop neuropsychiatric conditions, which are more prevalent in women than men. Efforts to model this sex difference in rodents have produced mixed results, with some studies paradoxically showing stronger phenotypes in males than females. Since changes in reproductive hormone levels can increase the likelihood of anxiety disorders in women, we examined the effects of ELA on adult female mice across the estrous cycle. We found that during diestrus, when the ratio of progesterone to estrogen is relatively high, ELA mice exhibit increased avoidance behavior, altered activity levels in specific contexts, and increased theta oscillation power in the ventral hippocampus. Ovariectomy, which eliminates circulating estrogen but not progesterone, unexpectedly preserved some of the effects present in diestrus ELA mice. Progesterone receptor antagonism in diestrus normalized avoidance behavior in ELA mice, while treatment with a negative allosteric modulator of the progesterone metabolite allopregnanolone promoted avoidance behavior in control mice. These results suggest that altered progesterone and allopregnanolone signaling during diestrus increases avoidance behavior in ELA mice

## Introduction

Early life adversity (ELA), which includes childhood maltreatment, chronic illnesses, accidents, natural disasters, and witnessing violence, significantly increases the likelihood of developing many forms of physical and mental illness in adulthood (Gallo et al., 2018; Li et al., 2016; Dunn et al., 2017; 2018). Among these conditions are anxiety disorders, the most prevalent psychiatric disorders (NIMH, 2018). Studies have shown that women are almost twice as likely to have an anxiety disorder diagnosis as men (NIMH, 2018; Somers et al., 2006; Craske and Stein, 2016) and that anxiety disorders in women are more disabling (McLean et al., 2011; GBD, 2017). While sex differences in response to ELA may not be evident during childhood, changes in hormone status are thought to “unmask” vulnerability (Hodes and Epperson, 2019). Indeed, some women experience excessive anxiety during times of dramatic reproductive hormone change, including puberty, pregnancy, childbirth, and menopause (both surgical and age-related) (Ross and Mclean, 2006; Vythilingum, 2008; Reardon et al., 2009; Flores-Ramos et al., 2018: Rocca et al., 2018), as well as at specific stages of the menstrual cycle (Christensen et al., 1992; Hantsoo and Epperson, 2015). Furthermore, the association between anxiety disorders and hormone change is more prevalent at each of these life stages/events for women who experienced childhood maltreatment (Choi and Sikkema, 2016; Zhu et al., 2019; Wakatsuki et al., 2020; Osofsky et al., 2021; Alexander et al., 2007; Shanmugan et al., 2020). Taken together, these findings suggest that ELA interacts with ovarian steroids to modulate vulnerability to anxiety disorders. However, the mechanisms that underlie this interaction remain unknown.

Efforts to understand how ELA affects the brain at the cellular and circuit levels in the service of vulnerability have involved the use of multiple animal models. Different mouse models of ELA have been shown to produce different behavioral phenotypes (Demaestri et al., 2020; Murthy and Gould, 2018; 2020), similar to human studies showing that different kinds of childhood maltreatment differentially predispose individuals to certain neuropsychiatric conditions (Huh et al., 2014; Infurna et al., 2016; Cohen et al., 2014). Operationalizing anxiety in the mouse can be problematic given the psychological aspects of anxiety in humans that involve conscious awareness (LeDoux and Pine, 2016). Less complex symptoms, such as avoidance or behavioral inhibition, as well as restlessness and agitation (Trombello et al., 2018; Franklin et al., 2021), may be effectively measured in mice by use of standard tests of avoidance behavior and locomotion, respectively. Using the ELA paradigm of maternal separation and early weaning (MSEW) in mice, we and others have found increased avoidance behavior and activity levels compared to control-reared mice (George et al., 2010; Carlyle et al., 2012; Murthy et al., 2019). These studies have either not tested female mice (George et al., 2010; Carlyle et al., 2012) or found no effect of MSEW on these behaviors in females (Murthy et al., 2019). The possibility that the estrous cycle may obscure effects of ELA has not yet been investigated.

Studies have shown that avoidance behavior and locomotion in mice, as well as self-reported anxiety in humans, are positively associated with neuronal oscillations in the theta range (4-12 Hz) in the hippocampus (McNaughton et al., 1983; Adhikari et al., 2010; Jacinto et al., 2013; Cornwell et al., 2012). In control mice, optogenetic stimulation of ventral hippocampus (vHIP) terminals in the medial prefrontal cortex at theta frequency increases avoidance behavior (Padilla-Coreano et al., 2019), and benzodiazepine treatment diminishes avoidance behavior coincident with decreased theta power (McNaughton and Gray, 2000; Yeung et al., 2012). Parvalbumin-positive (PV+) interneurons contribute to neuronal oscillations in the hippocampus by coordinating fast inhibition of principal neurons (Amilhon et al., 2015). A subpopulation of PV+ interneurons is surrounded by perineuronal nets (PNNs), specialized extracellular matrix structures that are known to regulate plasticity (Sorg et al., 2016). PNNs have been shown to alter neuronal oscillations (Carceller et al., 2020; Wingert and Sorg, 2021), raising the possibility that they are involved in the regulation of avoidance behavior. We have shown that MSEW increases PNNs surrounding vHIP PV+ cells and increases theta power coincident with increased avoidance behavior and activity levels (Murthy et al., 2019). However, these studies were not carried out in females, raising questions about whether similar mechanisms might underlie the connection between ELA and behavioral vulnerability.

To investigate whether ovarian status influences the effects of ELA on behavior, as well as on neuronal oscillations and PNNs in the vHIP, we examined control- and MSEW-reared female mice at different stages of estrous, including proestrus, estrus, metestrus, and diestrus, and after ovariectomy. We found that during diestrus, MSEW mice displayed increased avoidance behavior, reduced grooming, and altered locomotion in different contexts. Coincident with these behavioral effects, we observed increased theta power in the vHIP when MSEW mice were in diestrus, not estrus. In control mice, PNNs in the vHIP changed across the estrous cycle, an effect that was prevented in MSEW mice. However, MSEW-mediated changes in PNN size and composition were noted, but only during diestrus. Ovariectomy prevented several MSEW effects, but only in a low stress context, suggesting that ELA induces an underlying vulnerability that can be partially modulated by ovarian steroids.

Since ovariectomy eliminates circulating levels of estrogen but not progesterone, we next explored whether manipulating the activation of progesterone receptors or the action of the progesterone metabolite allopregnanolone would affect avoidance behavior. We found that progesterone receptor antagonism prevented the increase in avoidance behavior and reduction in grooming in MSEW diestrus mice, while inhibiting allopregnanolone action in control mice mimicked the increase in avoidance behavior of MSEW mice. Taken together with our observations that MSEW diestrus mice have decreased vCA1 expression of steroid 5α-reductase I, an enzyme involved in the local reduction of progesterone to its neurosteroid metabolites, these results suggest that diminished conversion of progesterone to allopregnanolone in ELA mice compared to control mice may contribute to increased avoidance behavior when mice are in diestrus.

## Results

### MSEW increases avoidance behavior during diestrus, but not estrus

To determine whether the effects of ELA on avoidance behavior are modulated by estrous cycle stage, we subjected mouse pups to the MSEW paradigm (Fig.1a,b), followed by testing on the elevated plus maze (EPM) in adulthood during proestrus, estrus, metestrus and diestrus. Since repeated testing on the EPM has been shown to change avoidance behavior (Schrader et al., 2018), we modified the task by increasing illumination and spraying a fine mist of water droplets on the open arms. In a pilot study, we tested mice on the “dry EPM” followed by two separate exposures to the “wet EPM”. We found that mice were significantly more avoidant of the open arms on the wet EPM than the dry EPM, and observed no evidence of habituation with repeated testing on the wet EPM (Fig. S1c; cycle (*F*_2,15_ = 0.3738, *p* = 0.0001, Tukey’s multiple comparisons test dry vs. wet 1 *p* = 0.0001, dry vs wet 2 *p* = 0.0001)). Therefore, we continued to use the wet EPM to assess avoidance behavior during each of the four stages of the estrous cycle (Fig.1c). When the data were combined across stages of the estrous cycle, no differences were observed in time spent or percent time spent in the open arms between control and MSEW mice (t_81_ = 0.9986, *p* = 0.3210) (Fig.1d). However, when the data were analyzed considering estrous stage as a variable (Fig.1e), a significant interaction was noted between estrous and ELA (Estrous x MSEW: *F*_3,55_ = 3.393, *p* = 0.0242), with a significant increase in avoidance behavior (i.e., a decrease in time spent and percent time spent on the open arms) between control and MSEW mice during diestrus (Sidak’s multiple comparisons test Control-MSEW Diestrus *p* = 0.0464) (Fig. 1f).

**Figure 1:**
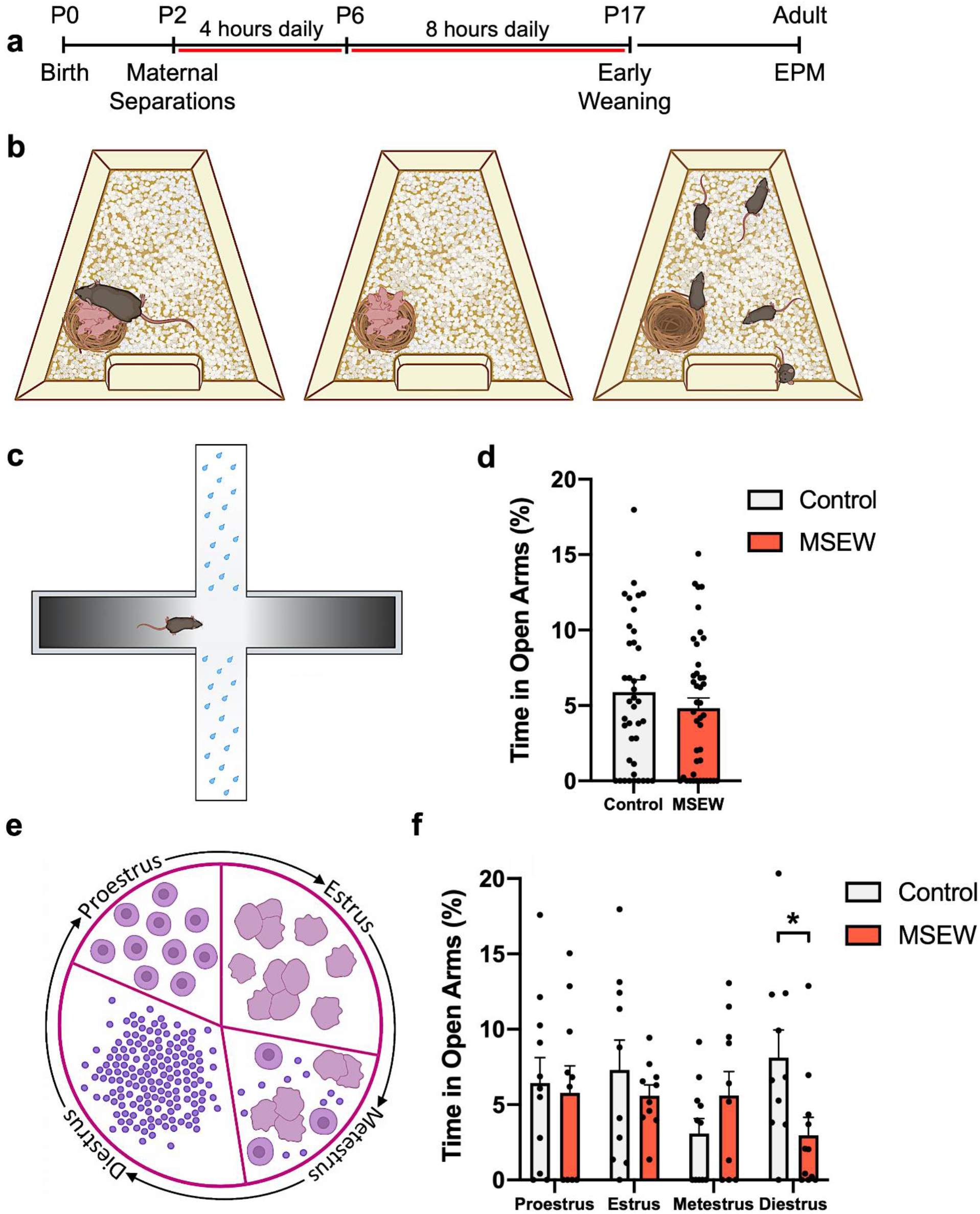
MSEW increases avoidance behavior only in diestrus. a,b) Timeline and schematics of MSEW paradigm. c) Schematic of the modified EPM, with water droplets on the open arms. d) Percent time in the open arms of the modified EPM is not significantly different between control and MSEW mice when stage of estrous during testing is not taken into consideration. e) Schematic of vaginal cytology used to determine whether mice were in different stages of the estrous cycle, proestrus, estrus, metestrus and diestrus. f) Testing on the modified EPM at specific stages of the estrous cycle shows that during diestrus, MSEW mice display increased avoidance behavior, i.e., reduced percent time in the open arms, compared to control mice. No significant differences between control and MSEW mice were observed during proestrus, estrus or metestrus. *p<0.05, two-way ANOVA (MSEW x Estrous followed by Sidak post hoc comparisons).

### MSEW alters activity levels in certain contexts during diestrus, but not estrus

We designed our subsequent studies to compare mice in estrus and diestrus, because these are the two longest estrous cycle phases (Ajayi and Akhigbe, 2020), with one phase, estrus, showing no significant difference in avoidance behavior between control and MSEW mice, while the other, diestrus, reveals more avoidance behavior in MSEW mice compared to control mice. We first observed behavior of control and MSEW mice in the home cage during estrus and diestrus and found no differences in activity levels (locomotion, climbing) between control and MSEW mice during estrus, but observed more locomotion (Estrous x MSEW: *F*_1,8_ = 7.132, *p* = 0.0283; Sidak’s multiple comparisons test control–MSEW Diestrus *p* = 0.0462) and a greater number of climbing bouts (Estrous *F*_1,12_ = 12.1, *p* = 0.0046, Sidak’s multiple comparisons test Estrus-Diestrus MSEW *p* = 0.0078) in MSEW diestrus mice only (Fig. 2c, d).

**Figure 2:**
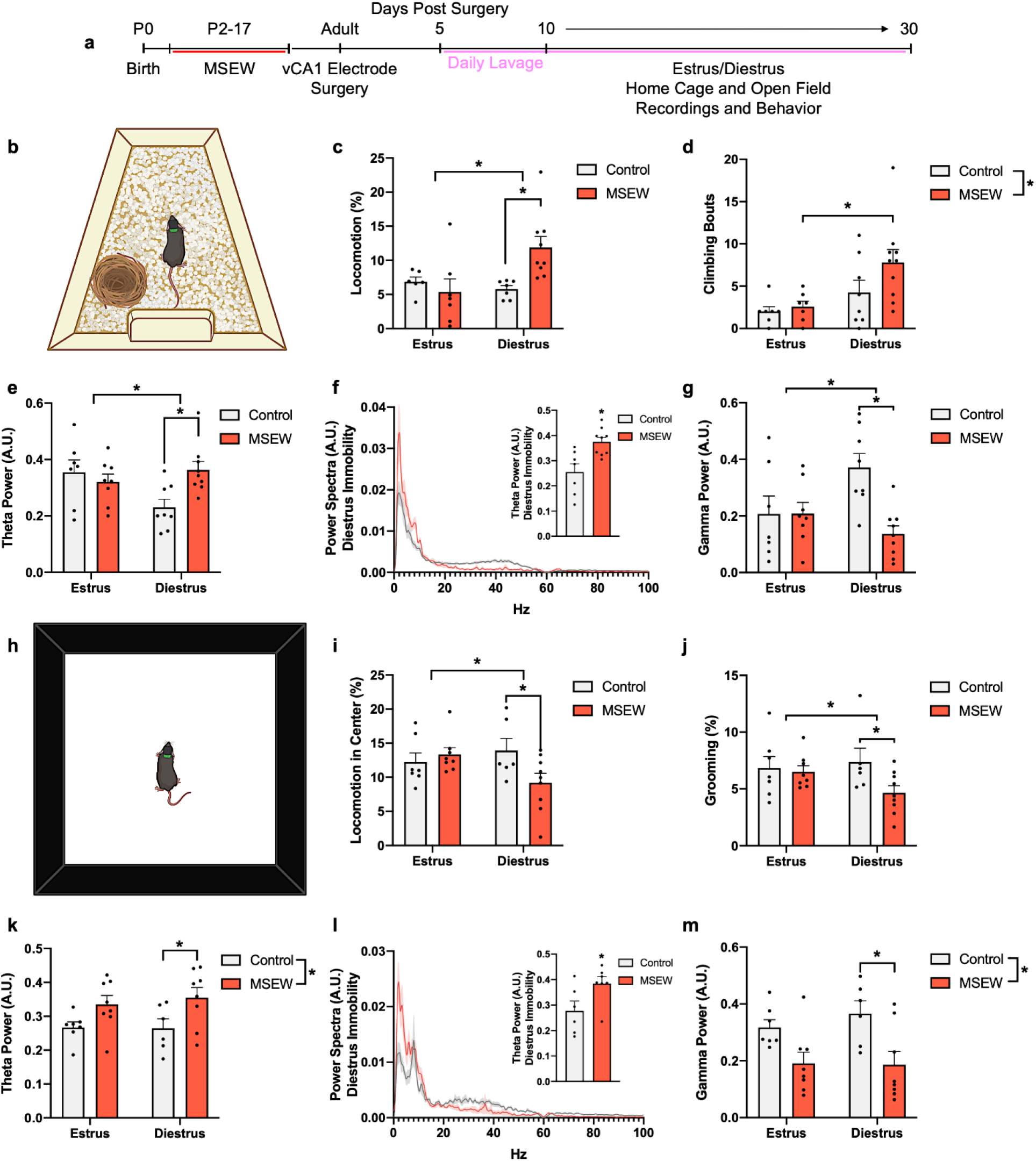
MSEW alters activity levels and vHIP theta oscillations depending on the context, and reduces grooming. a) Timeline of MSEW, surgery, lavage, behavioral testing and electrophysiology. b) Schematic of recording and testing in the home cage. c) Locomotion in the home cage is increased in MSEW mice in diestrus, but not estrus. d) Climbing in the home cage is increased in MSEW mice in diestrus, but not estrus. e) vHIP theta power is increased in MSEW mice in diestrus, but not estrus. f) Power spectra from diestrus vHIP LFPs during periods of immobility in the home cage from both control and MSEW mice, inset bar graph shows overall vHIP theta power is higher in diestrus MSEW mice during immobility compared to diestrus control mice. g) vHIP gamma power is higher in control diestrus compared to control estrus, but not in MSEW mice. h) Schematic of recording and testing in the open field. i) Locomotion in the center of the open field is reduced in MSEW mice during diestrus but not estrus. j) Grooming in the center of the open field is reduced in MSEW mice during diestrus but not estrus. k) vHIP theta power is significantly higher in the open field in MSEW mice during diestrus compared to control mice during diestrus, but not during estrus. l) Power spectra from diestrus vHIP LFPs during periods of immobility in the open field, inset bar graph shows overall vHIP theta power is higher in diestrus MSEW mice during immobility compared to diestrus control mice. m) vHIP gamma power is significantly reduced in MSEW mice compared to control mice in diestrus but not estrus. *<0.05, two-way ANOVA (MSEW x Estrous) followed by Sidak’s multiple comparison tests, or t tests for inset graphs (f, l).

We also assessed behavior in brightly-lit novel environments, which differed in translucence (clear and opaque) and lighting across trials to minimize habituation. We found no overall changes in locomotion between control and MSEW mice during estrus, but observed decreased locomotion in the center of the open field in MSEW mice during diestrus (Fig. 2i) and increased locomotion in the periphery of the open field in MSEW mice during diestrus (Estrous x MSEW: *F*_1,26_ = 4.489, *p* = 0.0438; Sidak’s multiple comparisons test control–MSEW diestrus *p* = 0.0462). We additionally measured another stress-sensitive behavior, grooming (Füzesi et la., 2016; Goodwill et al., 2019; Mu et al., 2020), and found no changes between control and MSEW mice while in estrus, but observed decreased grooming in MSEW mice during diestrus (Estrous x MSEW: *F*_1,11_ = 11.25, *p* = 0.0064, Sidak’s multiple comparison test control-MSEW Diestrus *p* = 0.0450) (Fig.2). Taken together, these findings suggest that MSEW diestrus mice display decreased locomotion in the center of the open field and decreased overall grooming, as well as increased activity when in contexts that are likely to be lower threat, i.e., the home cage and periphery of a novel environment.

*MSEW increases theta power in the ventral hippocampus during diestrus, but not estrus* We recorded LFPs from the ventral CA1 (vCA1) of control and MSEW female mice during behavioral testing, as vHIP theta power has been linked to avoidant behavior (Padilla-Coreano et al., 2019) and is increased in male mice after MSEW (Murthy et al., 2019). In the home cage, we observed significantly higher oscillatory power in the theta range (4-12 Hz) in MSEW mice while they were in diestrus, but not in estrus, compared to control mice (Estrous x MSEW: *F*_1,11_ = 6.872, *p* = 0.0238, Sidak’s multiple comparisons test Control-MSEW Diestrus *p* = 0.0115). Given that MSEW mice in diestrus displayed increased locomotion compared to control mice in diestrus, and because theta oscillations in the dorsal hippocampus (dHIP) have been linked to running (McNaughton et al., 1983), we further investigated whether theta power was increased only during periods of locomotion by analyzing LFPs during time-stamped behavioral epochs. MSEW mice demonstrated significantly higher vCA1 theta power in diestrus during periods of immobility than control mice (t_14_ = 3.384, *p* = 0.0045), and a correlational analysis of theta power in the vCA1 and locomotion revealed that although there is an overall group effect of increased locomotion and increased theta power in the MSEW diestrus group, increased locomotion is not driving the increase in theta power (Control: *r* = 0.04925, *p* = 0.8812, MSEW: *r* = 0.01204, *p* = 0.7787) (Fig. S1e).

In the open field, MSEW mice also showed an increase in vCA1 theta power compared to control mice during diestrus, but not estrus (Fig. 2k) (MSEW *F*_1,25_ = 9.068, *p* = 0.0059, Sidak’s multiple comparisons test Control-MSEW Diestrus *p* = 0.0498). This effect persisted throughout periods of immobility (t_11_ = 2.344, *p* = 0.0389) (Fig. 2i), and a correlational analysis of vCA1 theta power and open field locomotion revealed that increased locomotion does not drive increased theta power in the open field (Control: *r* = 0.2023, *p* = 0.3709, MSEW: *r* = 04870, *p* = 0.5683) (SFig. 1j). In both the home cage and open field, MSEW mice in diestrus displayed lower gamma oscillation power compared to control mice in diestrus (Estrous x MSEW: *F*_1,12_ = 10.63, *p* = 0.0068, Sidak’s multiple comparisons test Control-MSEW Diestrus *p* = 0.0013; MSEW *F*_1,14_ = 13.09, *p* = 0.0028, Sidak’s multiple comparisons test Control-MSEW Diestrus *p* = 0.0107). No significant change in gamma power was observed in MSEW mice during estrus (Fig. 2e, k).

### Ovariectomy has a context-dependent influence on behavior and ventral hippocampal theta power in MSEW mice

To determine whether differences in behavior and neuronal oscillations observed during diestrus in MSEW mice are dependent on ovarian steroids, control and MSEW mice were bilaterally ovariectomized (OVX) and, after recovery from surgery, tested in both conditions again. After OVX, MSEW mice spend more time moving in the open field (t_7_ = 2.509, *p* = 0.0405) compared to OVX control mice, and exhibit less grooming (t_7_ = 2.497, *p* = 0.0412). Coincident with increased movement, there was an increase in theta power in the vCA1 of OVX MSEW mice compared to OVX control mice (t_28_ = 2.224, *p* = 0.0344), alongside a decrease in gamma power (t_28_ = 3.284, *p* = 0.0027) (Fig.3, S3).

**Figure 3:**
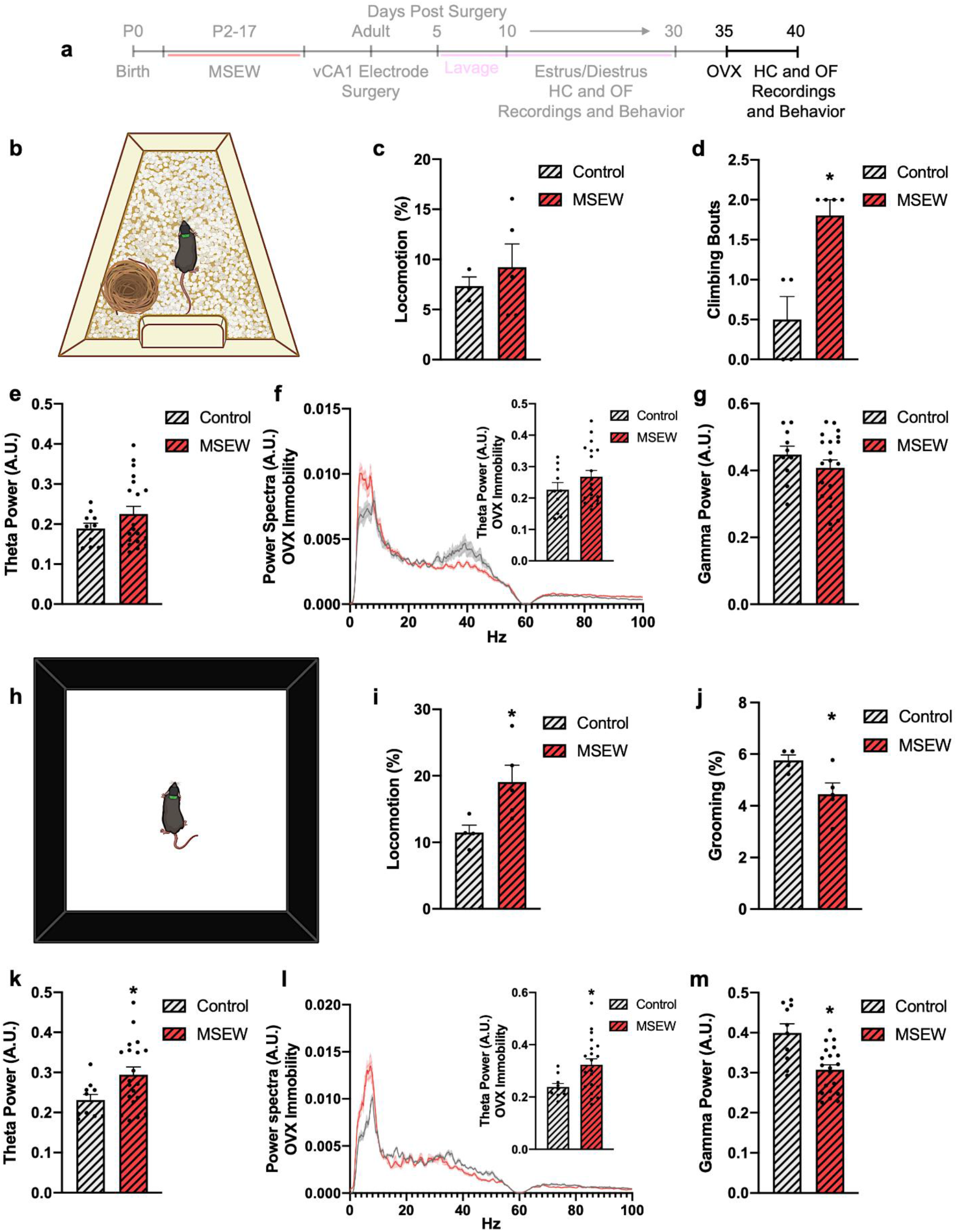
OVX reverses the effects of MSEW seen in diestrus mice, but only when they are in low stress contexts. a) Timeline of MSEW, electrode implantation, OVX, behavioral testing and electrophysiology. b) Schematic of recording and testing in the home cage. c) After OVX, locomotion in the home cage is not significantly different between control and MSEW. d) After OVX, MSEW mice still climb more than control mice. e) After OVX, vHIP theta power is not elevated in MSEW mice. f) Power spectra from OVX MSW and control mice in the home cage during periods of immobility; inset bar graph shows no significant difference in overall vHIP theta power between OVX control and MSEW. g) vHIP gamma power is no different between OVX MSEW and OVX control mice. h) Schematic of recording and testing in the open field. i) After OVX, locomotion in higher and j) grooming in the center of the open field is lower in MSEW compared to control. k) vHIP theta power is significantly higher in the open field in MSEW compared to control mice. l) Power spectra from OVX vHIP LFPs during periods of immobility in the open field; inset bar graph shows overall vHIP theta power is higher in OVX MSEW mice during immobility compared to OVX control mice. m) after OVX, vHIP gamma power is significantly reduced in MSEW mice compared to control mice. *<0.05, unpaired t tests.

Increased vCA1 theta power was observed during time-stamped bouts of immobility in OVX MSEW mice (t_28_ = 2.478, *p* = 0.0195), ruling out a locomotor-driven increase in theta power. By contrast, no significant increase in locomotion or theta power was observed in OVX MSEW mice in the home cage (Fig.3, S3). Collectively, these findings suggest that ovarian steroids are necessary for diestrus MSEW effects only under certain contexts; when OVX MSEW mice are in the familiar low-threat home cage environment, their behavior and neuronal oscillations are similar to OVX control mice, whereas in a novel, brightly-lit environment, locomotion and vCA1 theta oscillations are higher, similar to what is observed with MSEW diestrus mice.

### Estrous cycle mediated changes in ventral hippocampal PNNs are disrupted by MSEW and are partially reversed after OVX

PNNs surrounding PV+ interneurons have been linked to neuronal oscillations (Carceller et al., 2020; Wingert and Sorg, 2021) and are increased in the male mouse vHIP after MSEW (Murthy et al., 2019). Recent studies suggest that PNNs change alongside the diurnal rhythm (Pantazopoulos et al., 2020), raising the possibility that PNNs undergo additional change across the estrous cycle. To investigate this possibility, we perfused control and MSEW mice at the same time of day and examined PNNs across the estrous cycle.

In the ventral dentate gyrus (vDG), we found estrous cycle differences in the number of cells labeled with the lectin-based PNN marker *wisteria floribunda agglutinin* (WFA) (MSEW x Estrous: *F*_1,32_ = 14.26, *p* = 0.0007), with more cells observed during estrus than during diestrus (Sidak’s multiple comparisons Control-MSEW: Diestrus *p* = 0.0003) (Fig.4). A similar estrous cycle effect was observed when examining the number of PV+ cells with PNNs (Estrous: *F*_1,16_ = 8.200, *p* = 0.0113), although no differences were observed between diestrus and estrus, control or MSEW in the number of PV+ cells (Estrous: *F*_1,32_ = 3.025, *p* = 0.916) (Fig. S5). Because a subset of basket cells are CCK+, we examined this population and found no effects of estrous cycle or MSEW on cell density (Estrous: *F*_1,21_ = 2.272, *p* = 0.1168) (Fig. S5).

**Figure 4:**
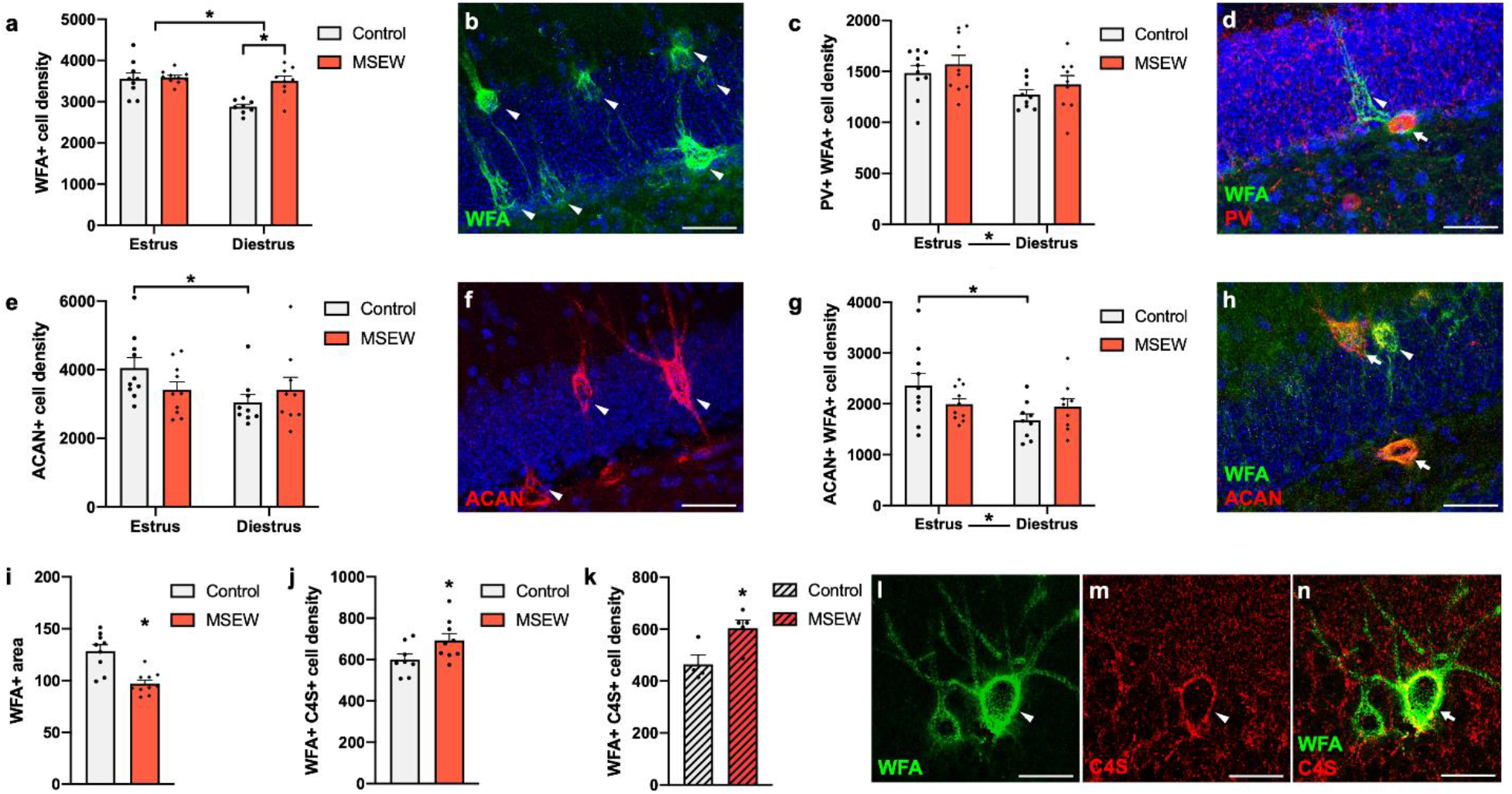
MSEW eliminates estrous cycle-mediated changes in the number of neurons with PNNs and alters PNN composition. a) The density of cells surrounded by WFA+ PNNs in the vDG is higher during estrus than diestrus in control mice; this difference does not exist in MSEW mice. b) Confocal example of WFA+ cells (green) in the vHIP. C) The density of PV+ cells surrounded by WFA+ PNNs in the vDG gyrus also changes across the estrous cycle. d) Confocal example of PV+ (red) WFA+ (green) cells in the vHIP. e) The density of cells surrounded by ACAN+ PNNs in the vDG gyrus is higher during estrus than diestrus in control mice; this difference does not exist in MSEW mice. f) Confocal example ACAN+ cells (red) in the vHIP. g) The density of ACAN+WFA+ cells in the vDG gyrus is higher in estrus than diestrus in control mice; this difference does not exist in MSEW mice. h) Confocal example of ACAN+ (green) WFA+ (red) cells in the vHIP. i) During diestrus, MSEW mice have lower PNN areas than control mice in ventral CA1. j) During diestrus, MSEW mice have a higher density of PNNs with C4S than control mice. k) After OVX, MSEW mice have higher density of WFA+ cells with C4S than control mice. l,m,o) Confocal images of two WFA+ (green) cells in the CA1, one of which is positive for C4S (red). *<0.05, two-way ANOVA (MSEW x Estrous) followed by Sidak’s multiple comparison tests, or t tests for MSEWvs control diestrus (i,j,k) and OVX (k).

To explore the influence of estrous cycle on PNNs in more depth, we also examined the main chondroitin sulfate proteoglycan (CSPG) aggrecan (ACAN). We found estrous cycle differences in ACAN+WFA+ cell density (Estrous: *F*_1,34_ = 4.817, *p* = 0.0344) with higher cell density in estrus compared to diestrus (Sidak’s multiple comparisons: *p* = 0.0130), but no estrous cycle differences in ACAN+ cell densities (Estrous: *F*_1,34_ = 3.035, *p* = 0.0905, Sidak’s multiple comparisons Estrus-Diestrus Control: *p* = 0.0377). In MSEW mice, estrous cycle differences in vDG PNNs were not evident in WFA+ cells (*p* = 0.7881), PV+WFA+ cells (*p* = 0.1523), ACAN+ cells (*p* = 0.999), or ACAN+WFA+cells (*p* = 0.9709) (Sidak’s multiple comparisons). In the vDG, no significant differences were observed in the percentage of PV+ cells that were WFA+, WFA+ cells that were PV+, ACAN+ cells that were WFA+, or WFA+ cells that were ACAN+ between estrous stages or control and MSEW (Table S1). These findings suggest that naturally occurring plasticity of PNNs surrounding PV+ cells across the estrous cycle is eliminated in MSEW mice. In the vCA1, estrous cycle or MSEW differences were not observed in the numbers or percentages of WFA+ (Estrous: *F*_1,34_ = 1.684, *p* = 0.2031), PV+WFA+ (Estrous: *F*_1,34_ = 0.009523, *p* = 0.9228), ACAN+WFA+ (Estrous: *F*_1,34_ = 0.3683, *p* = 0.5480), PV+ (Estrous: *F*_1,32_ = 0.3654, *p* = 0.5498), or CCK+ cells (Estrous: *F*_1,34_ = 0.5382, *p* = 0.4682) (Fig. S5; Table S2) However, vCA1 ACAN+ density counts revealed a significant interaction (Estrous x MSEW: *F*_1,34_ = 1.623, *p* = 0.0152).

Building off our findings that MSEW behavioral and vHIP oscillation effects are present only during diestrus, we investigated PNNs in control and MSEW diestrus mice in more detail. In the vCA1, we found differences in the size of PNNs surrounding PV+ cells. MSEW diestrus mice had smaller PNN cross sectional areas than estrus mice (t_16_ = 6.165, *p* = 0.0001), an effect that was also observed in the vDG (t_18_ = 3.641, *p* = 0.0019) (Fig. 4; S6). We observed no changes in PV+ area in the vDG (t_16_ = 01.026, *p* = 0.3203) or vCA1 (t_13_ = 0.4777, *p* = 0.9626).

Differences in PNN composition can influence neuronal function, leading us to examine the expression of the 4-sulfation pattern of chondroitin sulfate chains, which has been associated with reduced plasticity in PNNs (Yang et al., 2017; Foscarin et al., 2017). We found significant increases in C4S+WFA+ cells in the vCA1 (Fig. 4) (t_15_ = 2.178, *p* = 0.0458), but not the vDG (Fig. S6) (t_15_ = 0.4215, *p* = 0.6793), of MSEW diestrus mice compared to control diestrus mice. These findings indicate that in the vCA1, MSEW mice in diestrus have smaller PNNs with more PNNs containing C4S compared to control mice. After OVX, differences observed between control and MSEW diestrus mice were no longer evident for WFA+ cell density (t_7_ = 0.5536, *p* = 0.5971) or WFA+ area (t_7_ = 0.1673, *p* = 0.8719) (Fig. S6), but C4S+WFA+ cell density in the vCA1 remained higher in MSEW compared to control mice (t_7_ = 2.953, *p* = 0.0213) (Fig. 4).

### Progesterone and allopregnanolone signaling influences avoidance behavior in control and MSEW diestrus mice

We next sought to investigate whether progesterone and its metabolite allopregnanolone influence avoidance behavior. Diestrus mice were administered either vehicle, asoprisnil: a selective progesterone receptor modulator with primarily antagonistic action (DeManno et al., 2003), or sepranolone: a negative allosteric modulator of allopregnanolone’s GABA_A_R binding site (Bäckström et al., 2021), before being tested on the EPM and the open field. After vehicle administration, MSEW diestrus mice spent significantly less time in the open arms than control mice (Drug x MSEW: *F*_2,44_ = 13.00, *p* < 0.0001; Sidak’s multiple comparisons test control–MSEW vehicle *p* = 0.0163) (Fig. 5b). After asoprisnil administration, MSEW diestrus mice spent significantly more time in the open arms compared to their vehicle trial (Sidak’s multiple comparisons test MSEW vehicle-asoprisnil *p* = 0.0167), while control diestrus mice spent significantly less time in the open arms compared to their vehicle trial (Sidak’s multiple comparisons test control vehicle-asoprisnil *p* = 0.041) (Fig. 5b). Taken together, MSEW diestrus mice spent significantly more time in the open arms than control diestrus mice after asoprisnil administration (Sidak’s multiple comparisons test asoprisnil control–MSEW *p* = 0.0440) (Fig. 5b). Sepranolone administration eliminated the difference in time spent in the open arms between control and MSEW mice (Sidak’s multiple comparisons test sepranolone control–MSEW *p* = 0.7305) (Fig. 5b).

**Figure 5:**
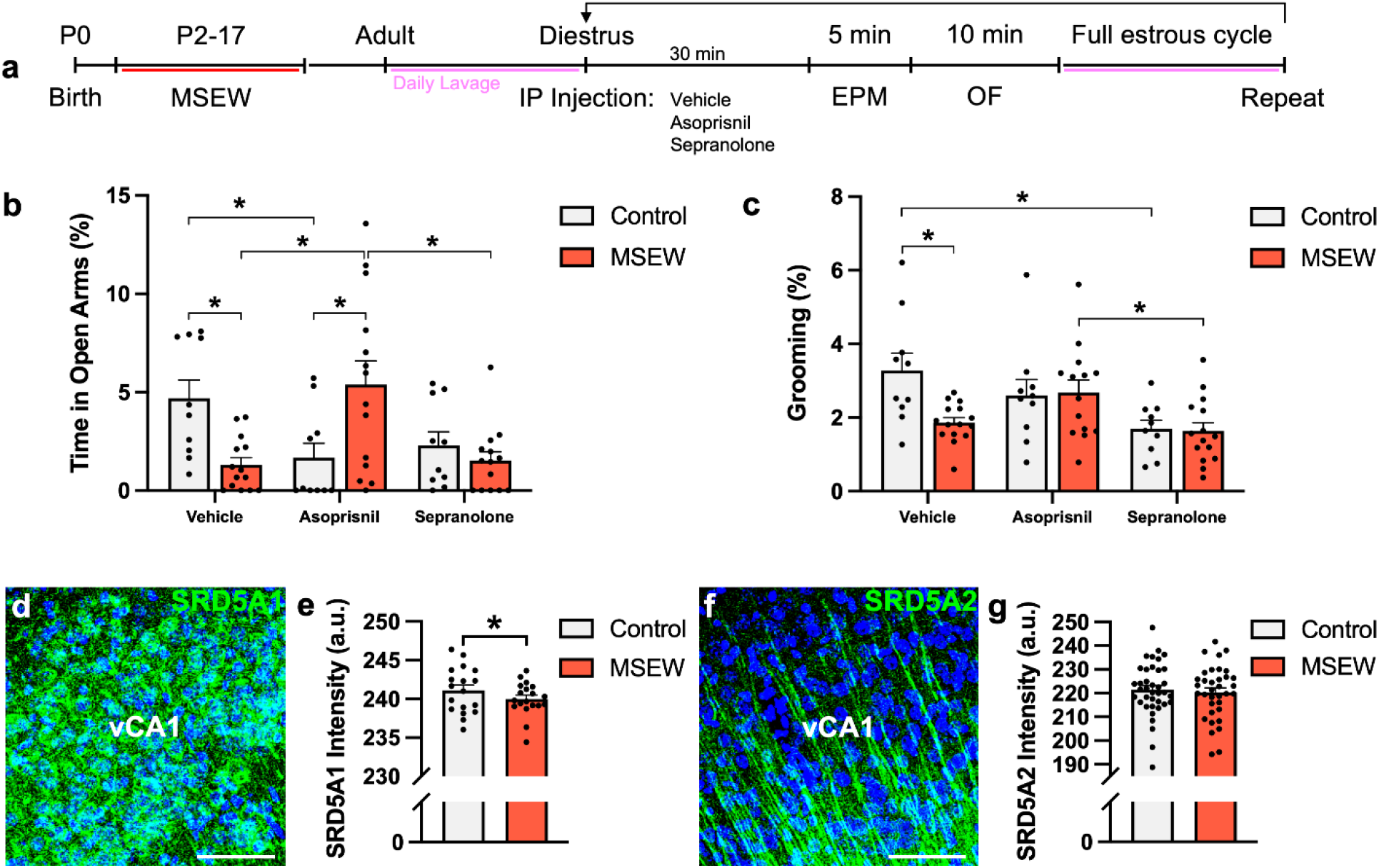
Progesterone receptor and allopregnanolone manipulations reverse control and MSEW diestrus differences in avoidance behavior. a) Timeline or behavioral paradigm. b) Percent time in the open arm after counterbalanced administration of vehicle, asoprisnil, and sepranolone in diestrus mice. Control mice spend more time in the open arm than MSEW mice after vehicle administration. MSEW mice undergo a significant increase in time spent in the open arms after asoprisnil administration. Control mice undergo a significant decrease in time spent in the open arms after asoprisnil administration. MSEW mice spend significantly more time in the open arms than control mice after asoprisnil administration. Control and MSEW mice have no significant difference in time spent in the open arms after separanolone administration. c) Percent time spent grooming while in the open field in diestrus mice. Control mice spend significantly more time grooming than MSEW mice after vehicle administration. Control mice spend significantly less time grooming after sepranolone administration compared to vehicle trials. d) Histology demonstrating average staining of SRD5A1 antibody in vCA1. e) MSEW diestrus mice exhibit significantly lower SRD5A1 intensity than control mice. f) Histology demonstrating average staining of SRD5A2 antibody in vCA1. g) Control and MSEW mice have no significant difference in vCA1 SRD5A2 intensity. *p<0.05, two-way ANOVA (MSEW x Drug followed by Sidak post hoc comparisons).

We next investigated grooming behavior in the open field after drug administration. Vehicle administration revealed that MSEW diestrus mice spent significantly less time grooming than control diestrus mice in the open field (Drug x MSEW: *F*_2,26_ = 5.722, *p* < 0.0063; Sidak’s multiple comparisons test vehicle control–MSEW *p* = 0.0068) (Fig. 5c). Administration of either asoprisnil or sepranolone abolished the difference in grooming time between control and MSEW mice (Sidak’s multiple comparisons test asoprisnil control-MSEW *p* = 0.9820; Sidak’s multiple comparisons test sepranolone control- MSEW *p* = 0.9988) (Fig. 5c). In control diestrus mice, sepranolone administration produced a significant decrease in time spent grooming when compared with their vehicle trial times (Sidak’s multiple comparisons test control vehicle-sepranolone *p* = 0.0044) (Fig. 5c).

Finally, we investigated the expression of steroid 5α-reductase type I and II in vCA1, the enzyme that facilitates the first of two steps in the metabolism of progesterone to allopregnanolone. A previous study reported reduced 5α-reductase expression in individuals who experienced ELA (Yehuda et al., 2009). We observed a significant decrease in vCA1 steroid 5α-reductase I intensity in diestrus MSEW mice compared to diestrus control mice (Linear mixed-effects regression *p* = 0.03017) (Fig. 5g). No difference was observed in steroid 5α-reductase II intensity (Linear mixed-effects regression *p* = 0.7438) (Fig. 5g).

## Discussion

Our findings suggest that the adverse effects of ELA may be unmasked in females during diestrus. We found that mice subjected to MSEW exhibited fluctuating levels of avoidance behavior on the EPM, with a significant increase in open arm avoidance during diestrus compared to control mice. No differences in avoidance behavior were observed between control and MSEW mice in other stages of the estrous cycle, including proestrus, estrus, or metestrus. We also found that during diestrus, MSEW mice showed increased activity levels in the home cage (locomotion, climbing) and increased locomotion in the periphery of the open field during diestrus, along with decreased locomotion in the center of the open field and decreased grooming, compared to control mice. Similar to what we observed for MSEW male mice (Murthy et al., 2019), behavioral differences between control and MSEW mice in diestrus were accompanied by increases in vHIP theta power in MSEW mice, including during periods of immobility, as well as alterations in vHIP PNNs. In contrast, none of these differences were observed between control and MSEW mice when they were in estrus.

These findings suggest an interaction between fluctuations in circulating ovarian hormones and ELA effects. To test this relationship, we eliminated the estrous cycle via ovariectomy and found that several of the behavioral and electrophysiological effects of MSEW persisted. While OVX induces floor levels of estrogen, progesterone production via the adrenal glands persists and is stress-sensitive, with stress elevating progesterone production (Romeo et al., 2004; Thorpe et al., 2014). The relatively high levels of progesterone might explain the persistence of diestrus-like behavior and vCA1 theta power in MSEW OVX mice when in the open field but not the home cage.

Previous studies have shown that PV+ interneurons play an important role in the generation of theta oscillations (Amilhon et al., 2015), and that these cells are altered by ELA in male mice (Murthy et al., 2019). A majority of PV+ interneurons are surrounded by PNNs, which are also influenced by ELA in the vDG of males (Murthy et al., 2019), and have been shown in other systems to affect neuronal oscillations (Carceller et al., 2020; Wingert and Sorg, 2021). Our findings suggest that PNNs change across the estrous cycle in control-reared mice and that this plasticity is disrupted after ELA. Estrous-mediated plasticity in PNNs may participate in buffering the hippocampus against adverse behavioral effects associated with fluctuations in ovarian steroids. In the absence of this plasticity after MSEW, times of a positive progesterone:estrogen ratio may result in increased avoidance behavior, altered activity levels, and reduced grooming. In addition to reduced estrous-mediated plasticity after MSEW, we observed reduced PNN size and an increased percentage of PNNs containing a chondroitin-4- sulfation pattern, a constituent associated with reduced plasticity (Yang et al., 2017; Foscarin et al., 2017), in the vCA1. The latter finding was also observed in OVX mice between control and MSEW groups, raising the possibility of potential causal links among C4S+ PNNs, increased theta oscillations, and alterations in behavior.

Because OVX eliminates circulating levels of estrogen but not progesterone, we next explored whether increased avoidance and decreased grooming behavior in MSEW diestrus mice could be altered by antagonizing progesterone receptors or inhibiting the action of the progesterone derivative allopregnanolone. We found that progesterone receptor antagonism prevented the effects of MSEW on avoidance and grooming behavior, while inhibiting allopregnanolone action in control mice mimicked the effects of MSEW. These findings raise the possibility that the conversion of progesterone to allopregnanolone is diminished in MSEW mice, an effect that is only functionally relevant during times of relatively high progesterone levels, e.g., during diestrus. In support of this hypothesis, we found that the progesterone metabolizing enzyme steroid 5α-reductase I is lower in the vHIP of MSEW mice.

While both estrus and diestrus are characterized by relatively low levels of circulating estrogen, progesterone levels rise during diestrus, yielding a higher progesterone:estrogen ratio than during estrus (McLean et al., 2012). Studies have shown that experimentally elevated progesterone levels can increase avoidance behavior in female mice by binding to progesterone receptors in the hippocampus (Galeeva and Tuohimaa, 2001; Galeeva et al., 2003; 2007), although many studies have reported that naturally occurring increases in progesterone levels across the estrous cycle do not have this effect (Plappert et al., 2005; Reynolds et al., 2018; Yohn et al., 2020). Previous studies suggest that ELA does not have a major impact on the estrous cycle or on circulating serum levels of estrogen in adult females (Eck et al., 2020; Manzano Nieves et al., 2019), raising the possibility that MSEW effects in diestrus may be primarily driven by different brain responses to changing levels of hormones instead of differences in circulating hormone levels themselves. These findings are consistent with human studies showing that excessive anxiety can emerge during times of ovarian steroid change, despite the fact that no clear correlations between anxiety and hormone levels exist (Hsiao et al., 2004: Azoulay et al., 2020; Hodes and Epperson, 2019).

In the healthy hippocampus, progesterone is metabolized to the neurosteroid allopregnanolone through two enzymes made by principal neurons (Agís-Balboa et al., 2007). Allopregnanolone is known to bind to GABA_A_ receptors in the hippocampus where it leads to reduced avoidance and defensive behaviors (Bitran et al., 1999; Rhodes and Frye, 2001; Smith et al., 2007; Modol et al., 2011). In the adult rodent brain, allopregnanolone levels and GABA_A_ receptor density are both modulated across the estrous cycle (Finn and Gee, 1993; Palumbo et al., 1995; Wu et al., 2013), with higher rates of conversion to allopregnanolone and binding of allopregnanolone to GABA_A_ receptors during diestrus than estrus (McCauley et al., 1995). These findings suggest that a buffering mechanism exists in the healthy brain to protect against potentially dysfunctional avoidance responses to natural increases in progesterone.

Studies have also shown that ELA reduces both allopregnanolone levels and GABA_A_ receptor binding (Frye et al., 2006; Mahmoodkhani et al., 2020), suggesting that this endogenous buffering mechanism may be disrupted after MSEW, although previous studies have not considered ELA-induced effects on these measures in the context of estrous stage. Our findings are consistent with the possibility that ELA-induced increases in avoidance behavior during diestrus are the result of diminished conversion of progesterone to allopregnanolone. We found that treatment with the selective progesterone receptor modulator asoprisnil blocked MSEW-induced increases in avoidance behavior, while treatment with sepranolone, an inhibitor of allopregnanolone action, increased avoidance behavior in control mice. Taken together with our findings of reduced 5α-reductase I expression in the vHIP of MSEW mice, the data suggest that elevated progesterone levels during diestrus produce increased avoidance behavior in MSEW mice due to an imbalance in the activation of progesterone receptors versus allopregnanolone (GABA_A_) receptors.

Studies have revealed that Holocaust survivors exhibit reduced 5α-reductase I expression, and that the most robust reductions were present in individuals that were youngest at the time of the war (Yehuda et al., 2009). Additional studies investigating postmortem brain tissue reveal that individuals with a major depression diagnosis exhibit diminished 5α-reductase I expression (Agis-Balboa et al., 2014). This effect was lost in individuals receiving antidepressant treatment at the time of death.

Human studies have shown that allopregnanolone is not only modulated across the menstrual cycle (Kimball et al., 2020), but is reduced in women with posttraumatic stress disorder (Pineles et al., 2018), a condition that is more prevalent in women who experienced childhood maltreatment (Lang et al., 2008). Along these lines, it is also worth noting that ELA predisposes women to premenstrual dysphoria, which often includes excessive anxiety (Azoulay et al., 2020).

Our findings suggest that theta power is elevated in MSEW mice during both diestrus and after OVX, coincident with behavioral effects suggesting altered stress-dependent activity levels and reduced grooming. Given that increased theta power was observed in MSEW mice when they are immobile, it is likely that the increased locomotion is not driving the increased theta, but the reverse may be the case. Further support comes from a number of studies demonstrating that increased theta power is only tightly coupled to locomotion speed in the dHIP and not the vHIP (Patel et al., 2012). vHIP theta power has been causally linked to increased avoidance behavior (Padilla-Coreano et al., 2019). Our findings suggest that this may be reflected in other behavioral effects, such as altered climbing and grooming. In this latter regard, it may be relevant that previous studies have shown an inverse correlation between grooming and theta power (Kemp and Kaada, 1975; Sainsbury et al., 1987; Dzirasa et al., 2010). Taken together, these findings suggest that increased vHIP theta power may be contributing to the increased avoidance behavior and the reduced locomotion and grooming in higher threat environments (e.g., the EPM and center of the open field), as well as the increased movement in low threat environments (e.g., the periphery of the open field and the home cage, perhaps akin to restlessness observed in humans with excessive anxiety) (Trombello et al., 2018; Franklin et al., 2021). It should be noted that a previous study suggested that ELA effects on grooming may be evidence of diminished “self-

care”, reflecting a “depressive-like” state (Goodwill et al., 2019). While it is not possible to know whether this was the case with the MSEW-induced diminished grooming we observed or whether it reflects behavioral inhibition in certain environments, it is likely relevant that there is a high comorbidity between major depressive disorder and anxiety disorders in humans (Kaiser et al., 2021).

## Methods

### Animals and MSEW paradigm

Animal procedures were approved by the Princeton University Institutional Animal Care and Use Committee and were in accordance with the National Research Council Guide for the Care and Use of Laboratory Animals (2011). Adult male and female C57BL/6J mice were obtained from The Jackson Laboratory and bred onsite at the Princeton Neuroscience Institute. On the day after birth, C57BL/6J pups were cross fostered and placed into control or MSEW litters. MSEW included maternal separation for 4 hours daily from P2-P5, maternal separation for 8 hours from P6-P16, and weaning at P17 (George et al., 2010; Murthy et al., 2019). Control litters were left undisturbed during this time and weaned at P21. During maternal separation, the dam was removed from the home cage and kept in the animal holding room in a clean cage with unlimited access to food and water. The home cage containing pups was moved to an adjacent room and placed on top of a heating pad maintained at 34° C. After weaning at P17 for MSEW mice and P21 for control mice, the groups of pups remained with their same sex littermates until behavioral testing in adulthood or surgery and behavioral testing in adulthood. Because the goal of this experiment was to explore effects of the estrous cycle on MSEW outcomes, only female offspring were used.

### Vaginal lavage and behavioral analyses

Control and MSEW mice were subjected to daily vaginal lavage beginning between 2-6 months of age to identify and track the estrous cycle of the mouse (Byers et al., 2012) before undergoing behavioral testing. Mice were only included in an estrous cycle study after they were observed to be cycling regularly through at least two cycles. For the first study, mice were examined in all four stages of the estrous cycle (proestrus, estrus, metestrus, diestrus). Thereafter, mice were selected for testing or perfusion when they were in estrus or diestrus.

### EPM testing

To measure avoidance behavior, the elevated plus maze (EPM) was used where avoidance of the open arms is considered to be evidence of avoidance behavior. Because repeated exposure to the EPM can result in habituation, we modified the testing apparatus to make the open arms more aversive by spraying them with water and increasing the brightness of the lamps (600 lux) over what we have typically used (200 lux; Murthy et al., 2019). Pilot studies in our lab showed that the wet EPM produces more avoidance of the open arms than the dry EPM, and does not show a change in behavior with repeated testing nor any change in entries to the closed arms (Fig. S1a-d). To avoid order effects, we counterbalanced exposure to the EPM across estrous cycle stages. On the day of testing, each mouse is placed into the center of the maze and their behavior was videotaped for 5 min. Time spent in open and closed arms, as well as number of entries into the arms, was determined by trained investigators watching coded videotapes so that the stage of estrous or ELA status remained unknown.

### Physical activity and stress-related behaviors

Locomotion and other stress-sensitive behaviors were measured in the home cage and two distinct open field boxes in separate groups of female mice with electrodes implanted in vCA1. The open field boxes differed in translucence (opaque and clear) and in overhead lighting (lit on one or two sides). Exposure to open field environments was counterbalanced across estrous stage to avoid habituation. Mice underwent ten-minute testing in each of the conditions when in estrus or diestrus. Locomotion was measured using scores of time spent engaged in locomotion, as well as the number of bouts of movement. The home cage had an elevated surface on one side and climbing times and bouts were recorded as an additional measure of physical activity. Grooming, a stress-sensitive behavior (Füzesi et la., 2016; Goodwill et al., 2019; Mu et al., 2020) was only measured in the open field, due to the low lighting of the home cage.

### Electrode implantation

Female control and MSEW mice were anesthetized and stereotaxically implanted with a custom-made 5 wire electrode array (Microprobes) into the unilateral vCA1 (AP: -3.5, ML: 3.4, DV: -3.5). After surgery, mice were singly housed to reduce overgrooming of the head stages. Behavioral testing and electrophysiological recording began two weeks after electrode implantation.

### Ovariectomy and progesterone/neurosteroid pharmacological manipulations

After testing in diestrus and estrus, electrode-implanted control and MSEW mice were subjected to bilateral ovariectomy (Strom et al., 2012). Mice were anesthetized and the ovaries located through a single midline incision on the dorsal surface. The uterine horn and vessels were ligated and the ovaries removed. Mice were allowed to recover for 5 days before electrophysiology and behavioral testing. Ovariectomy was confirmed by examining excised ovaries, and by examining the body cavity after perfusion.

Control and MSEW mice were administered vehicle, asoprisnil, or sepranolone 30 minutes prior to testing on the wet EPM and the open field. Drug administration was counterbalanced across mice. Mice spent five minutes on the EPM and 10 minutes in the open field. Testing on the wet EPM always occurred first and preceded open field testing by roughly 10 minutes At the conclusion of behavioral testing with a given drug, mice were excluded from additional testing until they had finished a complete estrous cycle and returned to diestrus.

### Electrophysiology

Local field potentials (LFPs) were recorded while mice were in the home cage and a brightly-lit open field using a wireless head stage (TBSI, Harvard Biosciences), in order to minimize stress during recording. Mice were tested during diestrus and estrus, with stage of estrous counterbalanced with order of testing in the novel environment and home cage. Control and MSEW mice were habituated to wearing the headstage in the home cage for 10 minutes a day for 5 consecutive days. After habituation, mice underwent LFP recordings during a 10 minute period in the home cage and a 10 minute period in a brightly-lit open field testing apparatus. LFPs were sent to a TBSI wireless 5 channel recording system, while mouse behavior was videotaped. The neural data were transmitted to a wireless receiver (Triangle Biosystems) and recorded using NeuroWare software (Triangle Biosystems). Continuous LFP data were highpass filtered at 1 Hz and notched at 60 Hz. All recordings referenced a silver wire wrapped around a ground screw implanted in the posterior parietal bone opposite of the electrode. Recordings were analyzed using Neuroexplorer software (Nex Technologies). To determine whether differences in neuronal oscillations were related to bouts of movement, separate analyses were done on mice during periods of locomotion and immobility. To normalize LFP data, the sum of power spectra values from 0 to 100 Hz were set to equal 1.

### Histology

Mice were anesthetized with Euthasol in estrus or diestrus and transcardially perfused with 4% paraformaldehyde. To avoid potential diurnal fluctuations in PNNs (Pantazopoulos et al., 2020), all mice were perfused at the same time of day.

Cryoprotected and frozen brain tissue was cut on a cryostat (Leica Biosystems) at 40 μm and a 1:6 series of sections was collected through the ventral hippocampus. Sections were pre-blocked in PBS containing 0.3% Triton X-100 and 3% normal donkey serum for 1.5 hr at room temperature. Sections were then incubated in biotin-conjugated *Wisteria floribunda* agglutinin (WFA, 1:1000, Sigma-Aldrich), mouse anti-PV (1:500, Sigma), rabbit anti-aggrecan (1:1000, Millipore), mouse anti-C6S (1:500, Sigma), mouse anti-C4S (1:500, Amsbio), rabbit anti-proCCK (1:500, Cosmo Bio), mouse anti-SRD5A1 (1:500, Proteintech), or rabbit anti-SRD5A2 (1:200, Invitrogen) for 24 h at 4°C, washed and then incubated in streptavidin Alexa Fluor 488 (1:1000, Invitrogen) or streptavidin Alexa Fluor 647 (1:1000), and secondary antibodies consisting of donkey anti-mouse Alexa Fluor 568 (1:500, Invitrogen), donkey anti-rabbit Alexa Fluor 488 (1:500 or 1:1000, Invitrogen), or donkey anti-mouse 568 (1:500, Invitrogen) for 1.5 hr at room temperature. Washed sections were then counterstained with Hoechst 33342 (1:5,000 Molecular Probes), mounted onto slides, and coverslipped over Vectashield (Vector Laboratories). See Table S3 for information about reagents.

Slides were coded until completion of the data analysis.

### Confocal microscopy and analyses

Z-stack images from vCA1 and vDG were taken with a Leica confocal microscope and the following analyses were carried out.

### Cell density measurements

For the analysis of cell densities during estrus and diestrus, and OVX, in control and MSEW brains, neuroanatomically-matched sections for each subregion were selected and the number of single and double labeled WFA and aggrecan stained cells, WFA and C4S double labeled cells, and single and double labeled WFA, PV or pro-CCK stained cells were counted on image stacks in ImageJ (NIH). Areas were measured using Image J software. Cell densities were determined for each mouse by dividing the total number of labeled cells by the volume for that subregion (area of subregion multiplied by 40 to account for thickness of cut section).

### Optical intensity, WFA and PV cell body area measurements

For the analysis of WFA and PV intensities, as well as PNN and cell body areas in control and MSEW diestrus brains, two neuroanatomically-matched sections were selected for each subregion for analysis. Z-stack 2μm optical images of WFA+ PV+ double labeled cells were analyzed in ImageJ (NIH). Prior to measuring intensity, the background was subtracted (rolling ball radius = 50 pixels). Using the ROI function, a perimeter was drawn around every individual WFA cell within that subregion, including the cell body and proximal dendrites. For each WFA cell, the maximum intensity value was calculated by multiplying the maximum mean gray value by the percent area. The maximum intensities of WFA and PV were averaged per section per animal.

### Optical intensity measurements

Z-stack images of the vCA1 were collected using a Leica SP8 confocal with LAS X software and a 40x oil objective. Settings remained constant throughout imaging. Collected SRD5A1 and SRD5A2 z-stack images were analyzed for optical intensity in Image J (NIH). A background subtraction using a rolling ball radius (50 pixels) was applied to the image stacks. A region of interest (ROI) was drawn around the pyramidal cell layer and the mean gray value was collected throughout the image stack.

### Statistical analyses

Behavioral data were collected and organized in Microsoft Excel. To analyze behavior and vCA1 LFP recordings in the home cage and open field, two-way ANOVA was used to compare data from control and MSEW mice in either estrus or diestrus. An unpaired t-t est was used to compare OVX behaviors and neuronal oscillation data in the home cage and open field. For OVX analyses, each electrode wire was included as a sample. Histochemical data were analyzed with unpaired t-tests for analyses that involved just two groups (control diestrus vs MSEW diestrus; control OVX vs MSEW OVX) or mixed-effects ANOVA for analyses that involved four groups (control diestrus, control estrus, MSEW diestrus, MSEW estrus). Data were presented as the mean + standard error of the mean (SEM). For all tests, a p-value of less than 0.05 was considered significant. Graphs were produced using GraphPad Prism 8.2.0.Acknowledgments: This work was supported by National Institute of Mental Health grant R01MH117459-01 (EG), a C.V. Starr Fellowship and a NARSAD Young Investigator Award (SSM).

## Acknowledgments

This work was supported by National Institute of Mental Health grant R01MH117459-01 (EG), a C.V. Starr Fellowship and a NARSAD Young Investigator Award (SSM).

---

**Figure S1:**
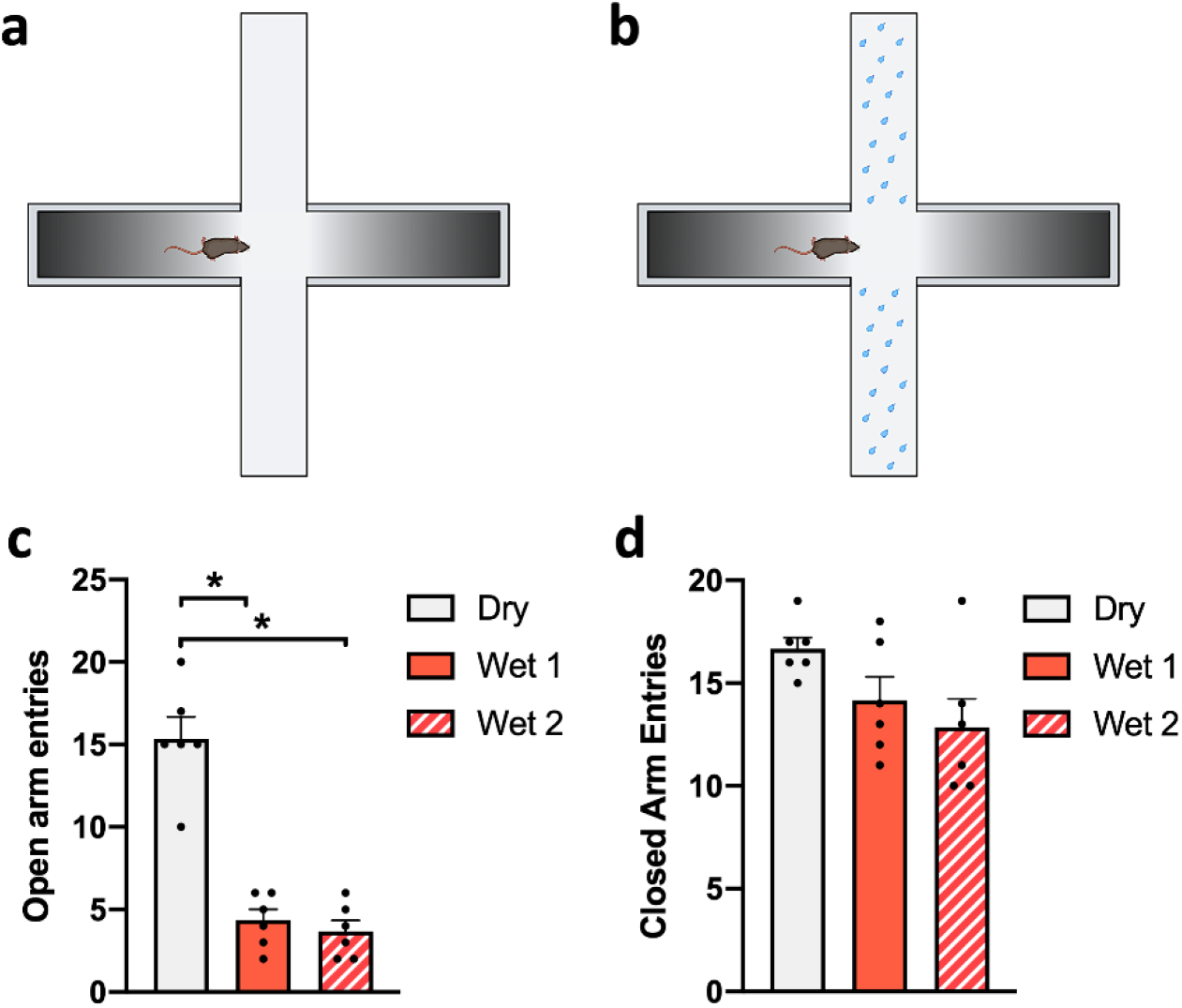
Avoidance behavior is increased on the modified “wet” EPM compared to the traditional “dry” EPM with no change after second exposure. Schematics of the traditional (a) and modified (b) EPM showing water droplets on open arms in b. c) Open arm entries are lower when exposed to the wet EPM compared to the dry EPM, with no difference between the first and second exposure to the wet EPM. d) Closed arm entries are not different when exposed to the dry or wet EPM. * p<0.05; repeated measures ANOVA with Tukey post hoc comparisons.

**Figure S2:**
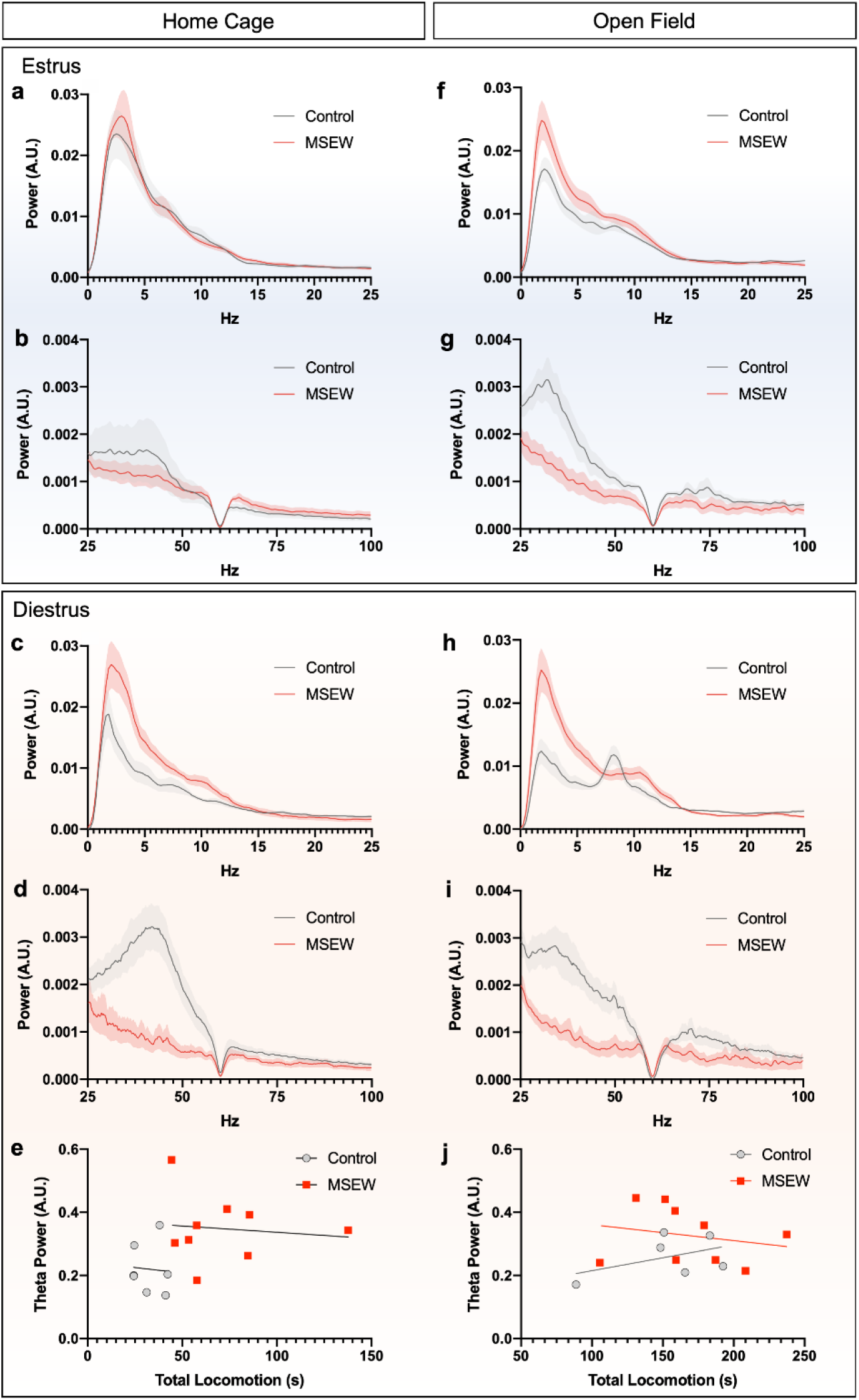
Estrous cycle MSEW and control vCA1 theta and gamma in the home cage and open field. **a, b** vCA1 power spectra in home cage during estrus. **c, d** vCA1 power spectra in home cage during diestrus. **e** Graph showing lack of correlation between theta power and locomotion duration in home cage diestrus animals. **f, g** vCA1 power spectra in open field during estrus. **h, i** vCA1 power spectra in open field during diestrus. **j** Graph showing lack of correlation between theta power and locomotion duration in open field diestrus animals.

**Figure S3:**
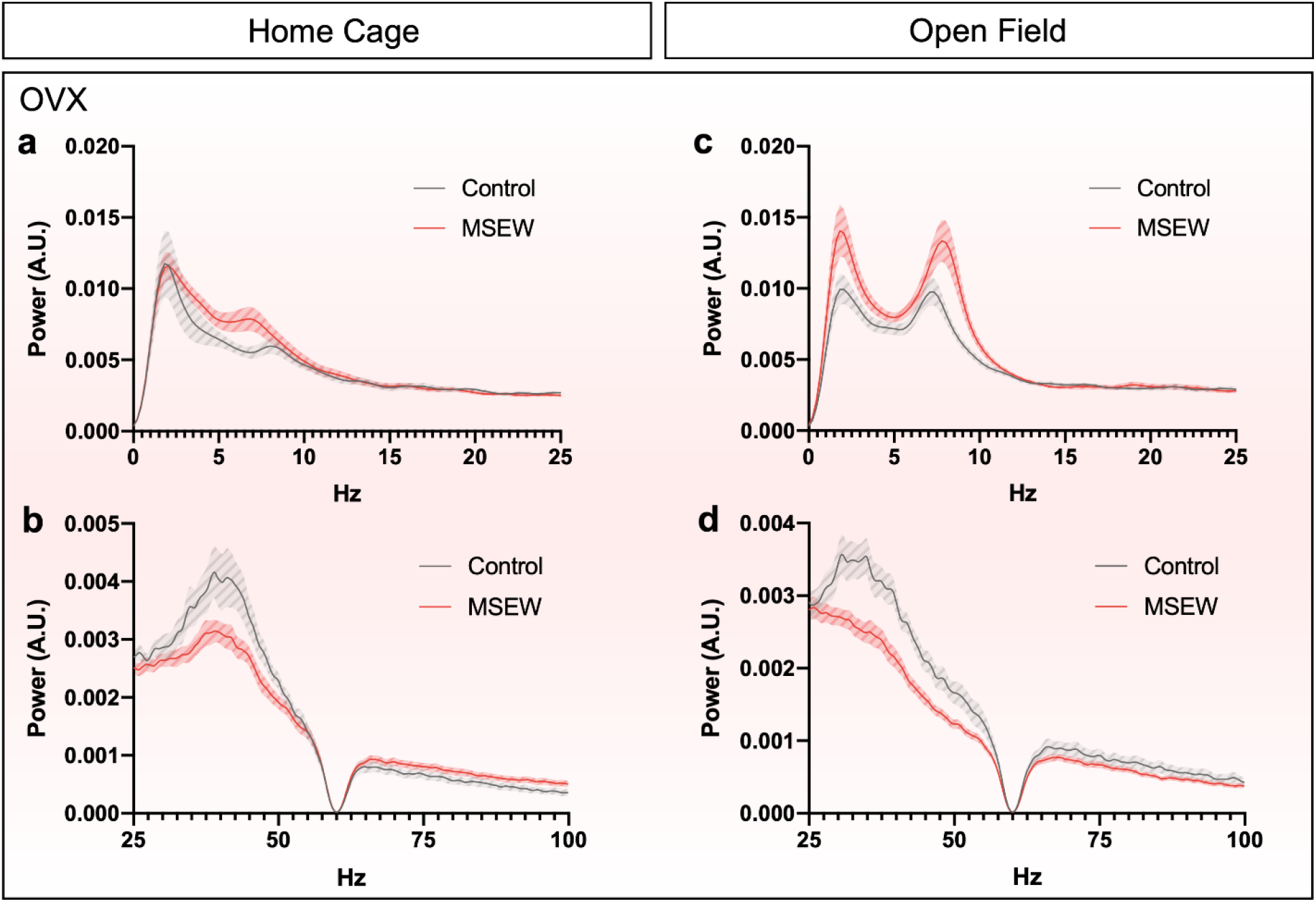
OVX MSEW and control vCA1 theta and gamma in the home cage and open field. **a, b** vCA1 power spectra in home cage after OVX. **c, d** vCA1 power spectra in open field after OVX.

**Figure S4:**
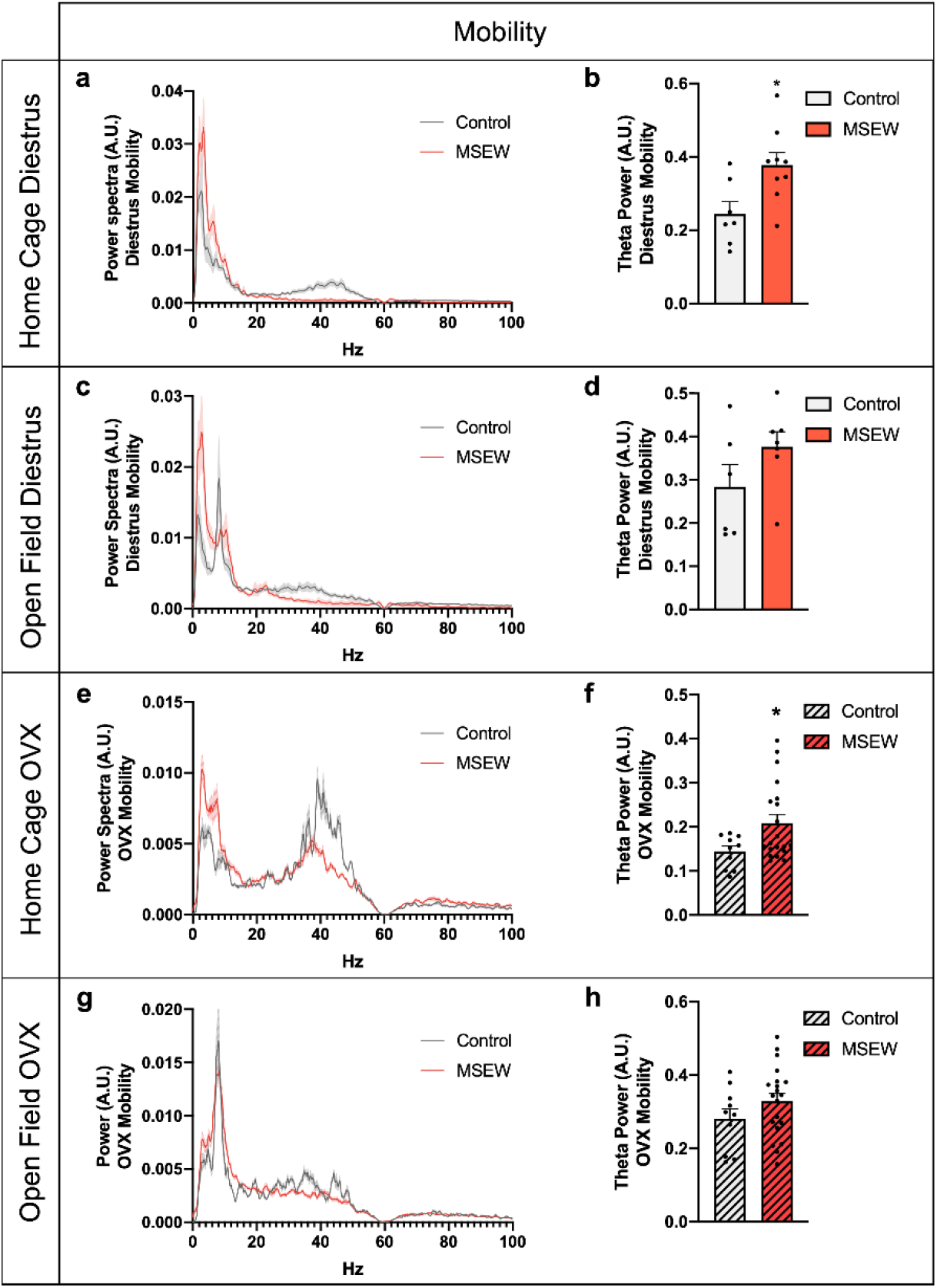
Estrous cycle and OVX MSEW and control vCA1 theta power in home cage and open field during mobility and immobility. **a** Diestrus vCA1 power spectra during mobility in home cage. **b** Diestrus vCA1 theta power during mobility in home cage. **c** Diestrus vCA1 power spectra during mobility in open field. **d** Diestrus vCA1 theta power during mobility in open field. **e** OVX vCA1 power spectra during mobility in home cage. **f** OVX vCA1 theta power during mobility in home cage. **g** OVX vCA1 power spectra during mobility in home cage. **h** OVX vCA1 theta power during mobility in home cage.

**Figure S5:**
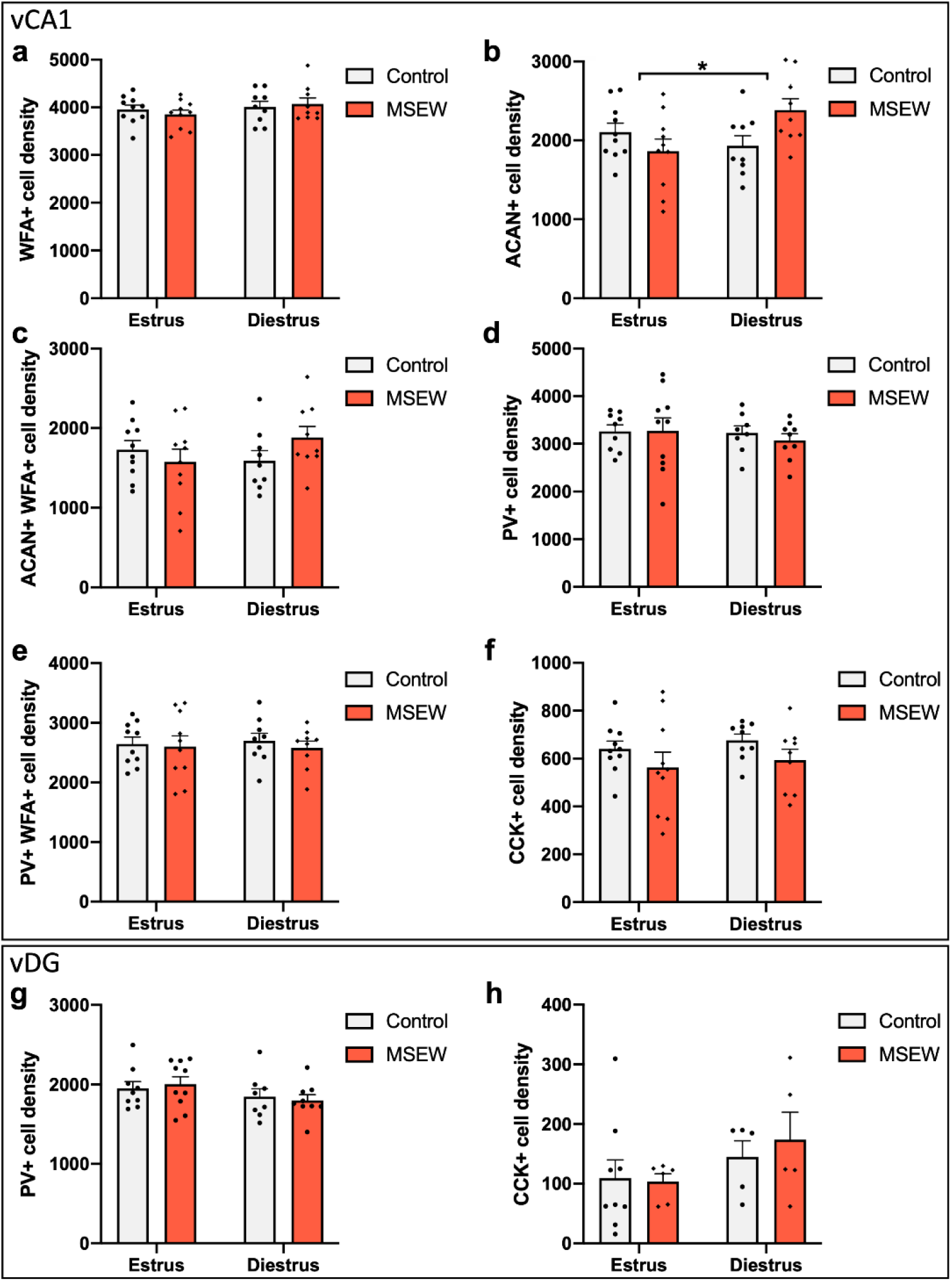
MSEW affects PNN size in diestrus. In the vCA1, no differences were detected in the density of WFA+ (a), PV+WFA+ (b), ACAN+WFA+ (d) or CCK+ cells (e) between estrus and diestrus in controls or MSEW. By contrast, mice in diestrus show more ACAN+ cells in MSEW compared to controls, but not in estrus (c). In the ventral dentate gyrus, no differences in CCK+ cells (f) or in WFA+ C4S+ cells (h) were detected across estrous stage in MSEW or control, although WFA+ PNN area is smaller in diestrus MSEW compared to controls. *<0.05, two-way ANOVA (MSEW x Estrous) followed by Sidak’s multiple comparison tests (a-f), or t tests for MSEW vs control diestrus (g,h).

**Figure S6:**
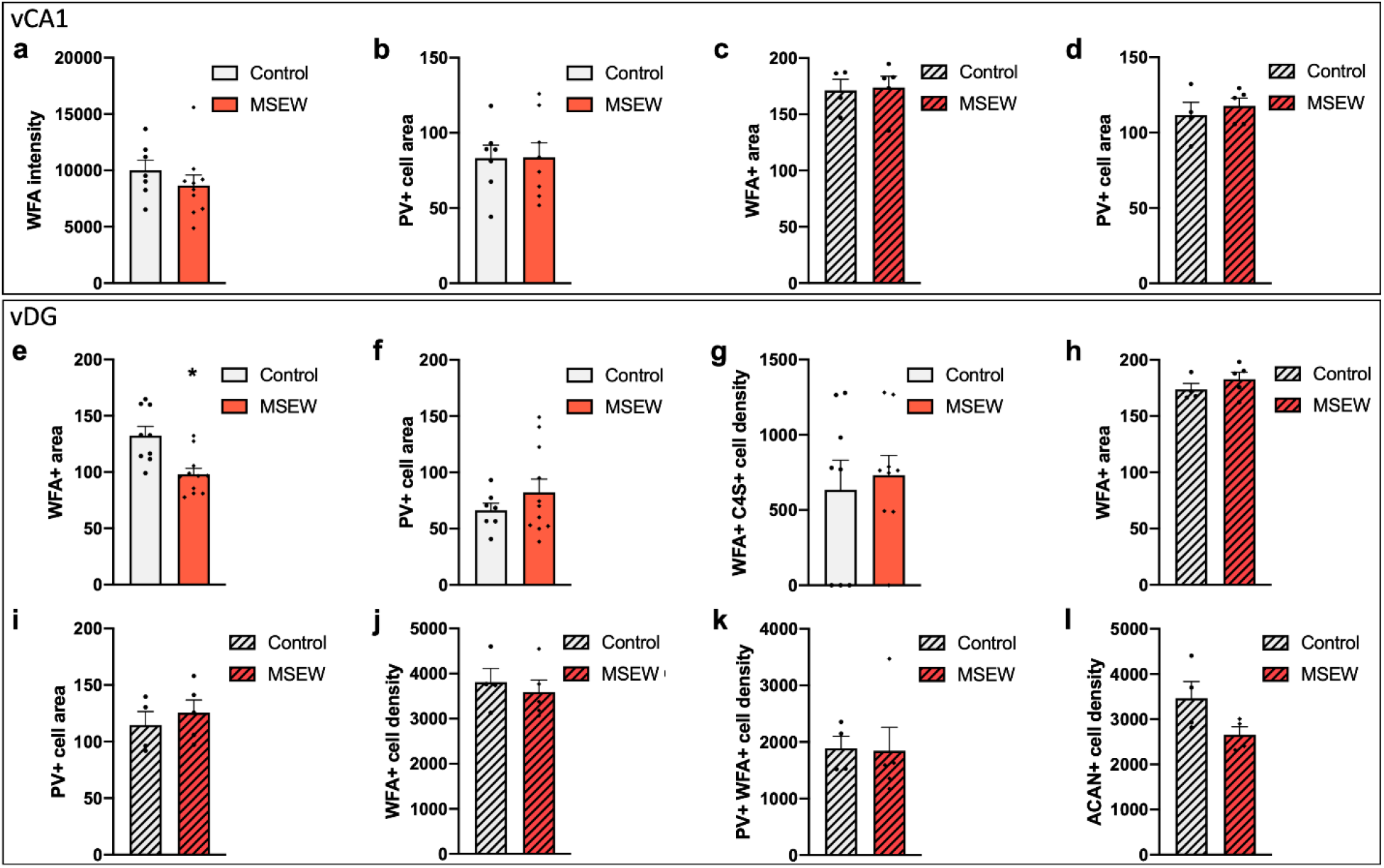
MSEW affects ventral dentate gyrus PNN size in diestrus. No differences were observed between control and MSEW during diestrus in WFA+ intensity (a), PV+ cell body area (b) or after OVX in WFA+ area (c) or PV+ cell body area (d). In the vDG, WFA+ area was smaller in diestrus MSEW compared to control mice (e). No differences in PV+ cell body area during diestrus (f) or WFA+C4S+ cell density during diestrus (g). After OVX, no differences were detected in WFA+ area (h), PV+ cell body area (i), WFA+ cell density (j), PV+WFA+ (k) or ACAN+ (l) cell density between control and MSEW. *<0.05, unpaired t tests for all panels.

**Table S1:** Percentage of double labeled cells in the ventral dentate gyrus. No significant differences were observed between control estrus and diestrus, or between control and MSEW within estrous stage or with OVX.

| vDG | Control Estrus | MSEW Estrus | Control Diestrus | MSEW Diestrus | Control OVX | MSEW OVX |
| --- | --- | --- | --- | --- | --- | --- |
| PV+WFA+/PV+ | 77.3±3.5 | 79.5±3.1 | 74.4±1.6 | 73.8±3.9 | 85.8±4.5 | 81.8±2.8 |
| PV+WFA+/WFA+ | 56.8±3.1 | 49.0±3.6 | 55.5±2.5 | 56.7±4.6 | 59.7±4.7 | 59.8±7.2 |
| ACAN+WFA+/ACAN+ | 57.9±2.5 | 60.0±2.4 | 55.6±2.3 | 58.1±3.3 | 62.9±2.0 | 69.4±5.6 |
| ACAN+WFA+/WFA+ | 53.9±2.3 | 52.2±2.9 | 45.8±2.0 | 46.6±2.7 | 63.8±1.1 | 65.8±4.5 |

**Table S2:** Percentage of double labeled cells in the ventral CA1 region

| vCAI | Control Estrus | MSEW Estrus | Control Diestrus | MSEW Diestrus | Control OVX | MSEW OVX |
| --- | --- | --- | --- | --- | --- | --- |
| PV+WFA+/PV+ | 82.0±1.1 | 76.2±3.5 | 82.7±2.3 | 83.9±1.2 | 84.1±3.1 | 82.3±4.1 |
| PV+WFA+/WFA+ | 74.0±1.7 | 67.2±7.5 | 72.4±1.5 | 71.8±2.4 | 58.1±4.8 | 54.9±2.7 |
| ACAN+WFA+/ACAN+ | 81.7±2.8 | 84.8±1.6 | 81.6±1.8 | 80.7±2.5 | 93.1±1.1 | 89.6±2.7 |
| ACAN+WFA+/WFA+ | 51.9±1.7 | 49.4±3.1 | 47.5±1.8 | 51.7±2.5 | 69.3±3.1 | 65.2±7.7 |

**Table S3:**
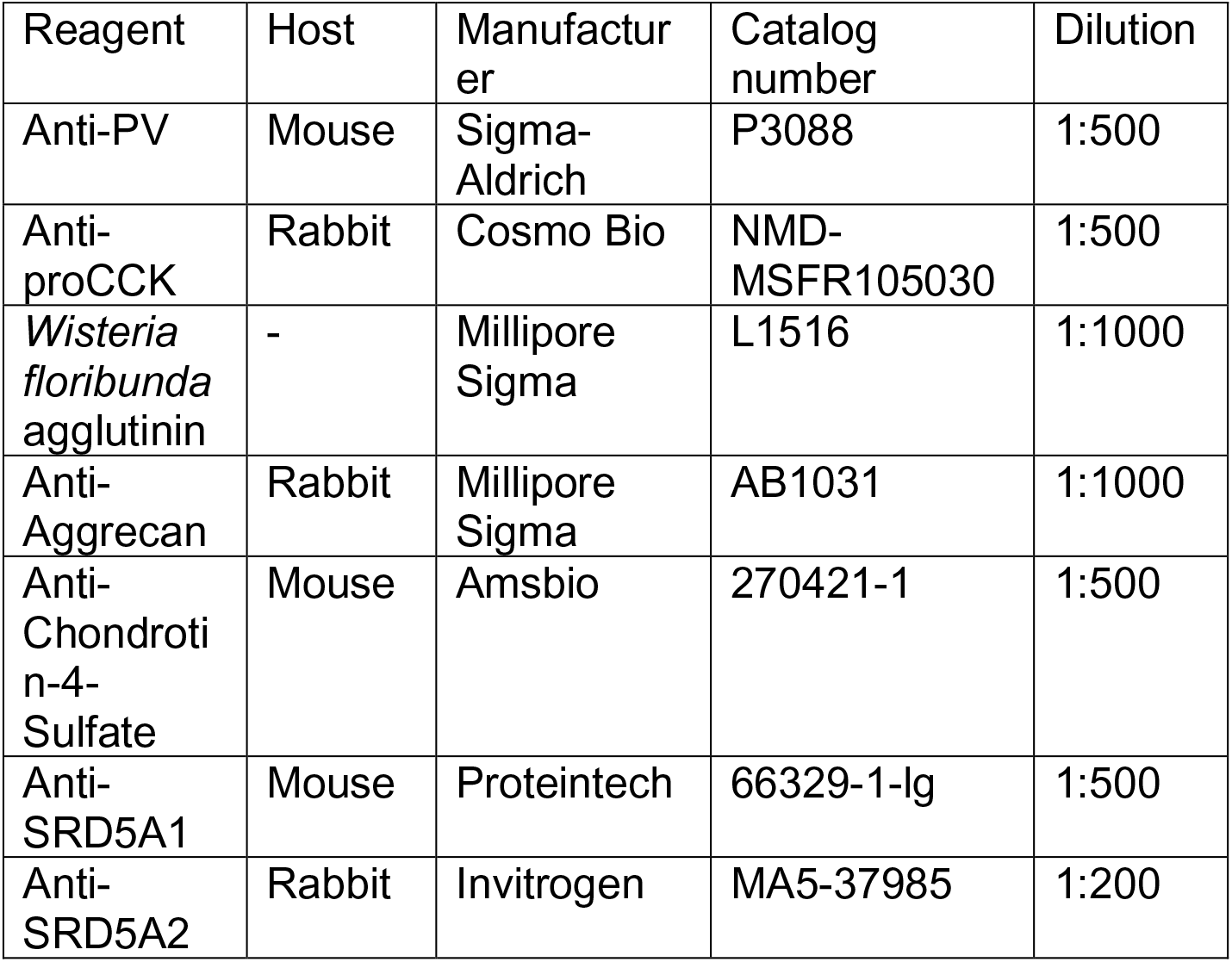
Reagents for Histochemistry

## Notes

### Competing Interest Statement

The authors have declared no competing interest.

## References

Adhikari A, Topiwala MA, Gordon JA (2010) Synchronized activity between the ventral hippocampus and the medial prefrontal cortex during anxiety. Neuron. 65(2):257–69.

Agis-Balboa RC, Guidotti A, Pinna G (2014) 5α-reductase type I expression is downregulated in the prefrontal cortex/Brodmann’s area 9 (BA9) of depressed patients. Psychopharmacology, 231(17): 3569–3580.

Agís-Balboa RC, Pinna G, Pibiri F, Kadriu B, Costa E, Guidotti A (2007) Down- regulation of neurosteroid biosynthesis in corticolimbic circuits mediates social isolation-induced behavior in mice. Proc Natl Acad Sci U S A. 104:18736–41.

Ajayi AF, Akhigbe RE (2020) Staging of the estrous cycle and induction of estrus in experimental rodents: an update. Fertil Res Pract. 14:6:5.

Alexander JL, Dennerstein L, Woods NF, McEwen BS, Halbreich U, Kotz K, Richardson G (2007) Role of stressful life events and menopausal stage in wellbeing and health. Expert Rev Neurother. 7(11 Suppl):S93–113.

Amilhon B, Huh CY, Manseau F, Ducharme G, Nichol H, Adamantidis A, Williams S (2015) Parvalbumin interneurons of hippocampus tune population activity at theta frequency. Neuron. 86(5):1277–89.

Azoulay M, Reuveni I, Dan R, Goelman G, Segman R, Kalla C, Bonne O, Canetti L. (2020) Childhood trauma and premenstrual symptoms: the role of emotion regulation. Child Abuse Negl. 108:104637.

Barrington J, Jarvis H, Redman JR, Armstrong SM (1993) Limited effect of three types of daily stress on rat free-running locomotor rhythms. Chronobiol Int. 10(6):410–9.

Bäckström T, Das R, Bixo M (2021). Positive GABA_A_ receptor modulating steroids and their antagonists: Implications for clinical treatments. Journal of neuroendocrinology, e13013.

Bitran D, Dugan M, Renda P, Ellis R, Foley M (1999) Anxiolytic effects of the neuroactive steroid pregnanolone (3 alpha-OH-5 beta-pregnan-20-one) after microinjection in the dorsal hippocampus and lateral septum. Brain Res. 850:217–24.

Byers SL, Wiles MV, Dunn SL, Taft RA (2012) Mouse estrous cycle identification tool and images. PLoS One. 7(4):e35538.

Carceller H, Guirado R, Ripolles-Campos E, Teruel-Marti V, Nacher J (2020) Perineuronal nets regulate the inhibitory perisomatic input onto parvalbumin interneurons and γ activity in the prefrontal cortex. J Neurosci. 40(26):5008–5018.

Carlyle BC, Duque A, Kitchen RR, Bordner KA, Coman D, Doolittle E, Papademetris X, Hyder F, Taylor JR, Simen AA (2012) Maternal separation with early weaning: a rodent model providing novel insights into neglect associated developmental deficits. Dev Psychopathol. 24(4):1401–16.

Choi KW, Sikkema KJ (2016) Childhood maltreatment and perinatal mood and anxiety disorders: a systematic review. Trauma Violence Abuse. 17(5):427–453.

Christensen AP, Board BJ, Oei TP (1992) A psychosocial profile of women with premenstrual dysphoria. J Affect Disord. 25:251–9.

Cohen LJ, Tanis T, Bhattacharjee R, Nesci C, Halmi W, Galynker I (2014) Are there differential relationships between different types of childhood maltreatment and different types of adult personality pathology? Psychiatry Res. 215(1):192–201.

Cornwell BR, Arkin N, Overstreet C, Carver FW, Grillon C (2012) Distinct contributions of human hippocampal theta to spatial cognition and anxiety. Hippocampus. 22(9):1848–59.

Demaestri C, Pan T, Critz M, Ofray D, Gallo M, Bath KG (2020) Type of early life adversity confers differential, sex-dependent effects on early maturational milestones in mice. Horm Behav. 124:104763.

DeManno D, Elger W, Garg R, Lee R, Schneider B, Hess-Stumpp H, Schubert G, and Chwalisz K (2003) Asoprisnil (J867): a selective progesterone receptor modulator for gynecological therapy. Steroids, 68(10-13): 1019–1032.

Craske MG, Stein MB (2016) Anxiety. Lancet. 388(10063):3048–3059.

Dunn EC, Nishimi K, Gomez SH, Powers A, Bradley B (2018) Developmental timing of trauma exposure and emotion dysregulation in adulthood: Are there sensitive periods when trauma is most harmful? J Affect Disord. 227:869–877.

Dunn EC, Nishimi K, Powers A, Bradley B (2017) Is developmental timing of trauma exposure associated with depressive and post-traumatic stress disorder symptoms in adulthood? J Psychiatr Res. 84:119–127.

Dzirasa K, Phillips HW, Sotnikova TD, Salahpour A, Kumar S, Gainetdinov RR, Caron MG, Nicolelis MA (2010) Noradrenergic control of cortico-striato-thalamic and mesolimbic cross-structural synchrony. J Neurosci. 30(18):6387–97.

Eck SR, Ardekani CS, Salvatore M, Luz S, Kim ED, Rogers CM, Hall A, Lee DE, Famularo ST, Bhatnagar S, Bangasser DA (2020) The effects of early life adversity on growth, maturation, and steroid hormones in male and female rats. Eur J Neurosci. 52(1):2664–2680.

Finn DA, Gee KW (1993) The influence of estrus cycle on neurosteroid potency at the gamma-aminobutyric acidA receptor complex. J Pharmacol Exp Ther. 265:1374–9.

Flores-Ramos M, Silvestri Tomassoni R, Guerrero-López JB, Salinas M (2018) Evaluation of trait and state anxiety levels in a group of peri- and postmenopausal women. Women Health. 58(3):305–319.

Foscarin S, Raha-Chowdhury R, Fawcett JW, Kwok JCF (2017) Brain ageing changes proteoglycan sulfation, rendering perineuronal nets more inhibitory. Aging (Albany NY). 9(6):1607–1622.

Franklin AR, Mathersul DC, Raine A, Ruscio AM (2021) Restlessness in generalized anxiety disorder: using actigraphy to measure physiological reactions to threat. Behav Ther. 52(3):734–744.

Frye CA, Rhodes ME, Raol YH, Brooks-Kayal AR. (2006) Early postnatal stimulation alters pregnane neurosteroids in the hippocampus. Psychopharmacology. 186:343–50.

Füzesi T, Daviu N, Wamsteeker Cusulin JI, Bonin RP, Bains JS (2016) Hypothalamic CRH neurons orchestrate complex behaviours after stress. Nat Commun. 7:11937

Galeeva A, Tuohimaa P. (2001) Analysis of mouse plus-maze behavior modulated by ovarian steroids. Behav Brain Res. 119(1):41–7.

Galeeva AY, Tuohimaa P, Shalyapina VG (2003) The role of sex steroids in forming anxiety states in female mice. Neurosci Behav Physiol. 33:415–20.

Galeeva AY, Pivina SG, Tuohimaa P, Ordyan NE. (2007) Involvement of nuclear progesterone receptors in the formation of anxiety in female mice. Neurosci Behav Physiol.37:843–8.

Gallo EAG, De Mola CL, Wehrmeister F, Gonçalves H, Kieling C, Murray J (2017) Childhood maltreatment preceding depressive disorder at age 18 years: A prospective Brazilian birth cohort study. J Affect Disord. 217:218–224.

GBD (2017) Disease and Injury Incidence and Prevalence Collaborators. Global, regional, and national incidence, prevalence, and years lived with disability for 328 diseases and injuries for 195 countries, 1990-2016: a systematic analysis for the Global Burden of Disease Study Lancet. 390(10100):1211–1259.

George, E.D., Bordner, K.A., Elwafi, H.M. et al. (2010) Maternal separation with early weaning: a novel mouse model of early life neglect. BMC Neurosci 11, 123.

Goodwill HL, Manzano-Nieves G, Gallo M, Lee HI, Oyerinde E, Serre T, Bath KG (2019) Early life stress leads to sex differences in development of depressive-like outcomes in a mouse model. Neuropsychopharmacology. 44:711–720.

Hantsoo L, Epperson CN (2015) Premenstrual dysphoric disorder: epidemiology and treatment. Curr Psychiatry Rep. 17(11):87.

Hodes GE, Epperson CN (2019) Sex differences in vulnerability and resilience to stress across the life span. Biol Psychiatry. 86(6):421–432.

Hsiao CC, Liu CY, Hsiao MC (2004) No correlation of depression and anxiety to plasma estrogen and progesterone levels in patients with premenstrual dysphoric disorder. Psychiatry Clin Neurosci. 58:593–9.

Huh HJ, Kim SY, Yu JJ, Chae JH (2014) Childhood trauma and adult interpersonal relationship problems in patients with depression and anxiety disorders. Ann Gen Psychiatry. 16;13:26

Infurna MR, Reichl C, Parzer P, Schimmenti A, Bifulco A, Kaess M (2016) Associations between depression and specific childhood experiences of abuse and neglect: A meta-analysis. J Affect Disord. 190:47–55.

Jacinto LR, Reis JS, Dias NS, Cerqueira JJ, Correia JH, Sousa N (2013) Stress affects theta activity in limbic networks and impairs novelty-induced exploration and familiarization. Front Behav Neurosci. 7:127.

Kaiser T, Herzog P, Voderholzer U, Brakemeier EL (2021) Unraveling the comorbidity of depression and anxiety in a large inpatient sample: Network analysis to examine bridge symptoms. Depress Anxiety. 38:307–317.

Kemp IR, Kaada BR (1975) The relation of hippocampal theta activity to arousal, attentive behaviour and somato-motor movements in unrestrained cats. Brain Res. 95(2-3):323–42.

Kimball A, Dichtel LE, Nyer MB, Mischoulon D, Fisher LB, Cusin C, Dording CM, Trinh NH, Yeung A, Haines MS, Sung JC, Pinna G, Rasmusson AM, Carpenter LL, Fava M, Klibanski A, Miller KK (2020) The allopregnanolone to progesterone ratio across the menstrual cycle and in menopause. Psychoneuroendocrinology. 112:104512.

Lang AJ, Aarons GA, Gearity J, Laffaye C, Satz L, Dresselhaus TR, Stein MB (2008) Direct and indirect links between childhood maltreatment, posttraumatic stress disorder, and women’s health. Behav Med. 33(4):125–35.

Li M, D’Arcy C, Meng X (2016) Maltreatment in childhood substantially increases the risk of adult depression and anxiety in prospective cohort studies: systematic review, meta-analysis, and proportional attributable fractions. Psychol Med. 46(4):717–30.

LeDoux JE, Pine DS (2016) Using neuroscience to help understand fear and anxiety: a two-system framework. Am J Psychiatry.173(11):1083–1093.

Mahmoodkhani M, Ghasemi M, Derafshpour L, Amini M, Mehranfard N (2020) Long-term decreases in the expression of calcineurin and GABAa receptors induced by early maternal separation are associated with increased anxiety-like behavior in adult male rats. Dev Neurosci. 42(2-4):135–144.

Manzano Nieves G, Schilit Nitenson A, Lee HI, Gallo M, Aguilar Z, Johnsen A, Bravo M, Bath KG (2019) Early life stress delays sexual maturation in female mice. Front Mol Neurosci. 12:27.

McCauley LD, Gee KW (1995) Influence of the estrus cycle on the discrimination of apparent neuroactive steroid site subtypes on the gamma-aminobutyric acidA receptor complex in the rat. J Pharmacol Exp Ther. 275(3):1412–7.

McLean CP, Asnaani A, Litz BT, Hofmann SG (2011) Gender differences in anxiety disorders: prevalence, course of illness, comorbidity and burden of illness. J Psychiatr Res. 45(8):1027–35.

McLean AC, Valenzuela N, Fai S, Bennett SA (2012) Performing vaginal lavage, crystal violet staining, and vaginal cytological evaluation for mouse estrous cycle staging identification. J Vis Exp. 67:e4389.

McNaughton N, Gray JA (2000) Anxiolytic action on the behavioural inhibition system implies multiple types of arousal contribute to anxiety. J Affect Disord. 61(3):161–76.

Mòdol L, Darbra S, Pallarès M (2011) Neurosteroids infusion into the CA1 hippocampal region on exploration, anxiety-like behaviour and aversive learning. Behav Brain Res. 222(1):223–9.

Mu MD, Geng HY, Rong KL, Peng RC, Wang ST, Geng LT, Qian ZM, Yung WH, Ke Y (2020) A limbic circuitry involved in emotional stress-induced grooming. Nat Commun. 11(1):2261.

Murthy S, Gould E (2020) How early life adversity influences defensive circuitry. Trends Neurosci. 43:200–212.

Murthy S, Kane GA, Katchur NJ, Lara Mejia PS, Obiofuma G, Buschman TJ, McEwen BS, Gould E (2019) Perineuronal nets, inhibitory interneurons, and anxiety-related ventral hippocampal neuronal oscillations are altered by early life adversity. Biol Psychiatry. 85(12):1011–1020.

NIMH (2018) U.S. Department of Health and Human Services. Anxiety disorders. National Institute of Mental Health. https://www.nimh.nih.gov/health/topics/anxiety-disorders.

Osofsky JD, Osofsky HJ, Frazer AL, Fields-Olivieri MA, Many M, Selby M, Holman S, Conrad E. (2021) The importance of adverse childhood experiences during the perinatal period. Am Psychol. 76(2):350–363.

Padilla-Coreano N, Canetta S, Mikofsky RM, Alway E, Passecker J, Myroshnychenko MV, Garcia-Garcia AL, Warren R, Teboul E, Blackman DR, Morton MP, Hupalo S, Tye KM, Kellendonk C, Kupferschmidt DA, Gordon JA (2019) Hippocampal-prefrontal theta transmission regulates avoidance behavior. Neuron. 104(3):601–610.e4.

Palumbo MA, Salvestroni C, Gallo R, Guo AL, Genazzani AD, Artini PG, Petraglia F, Genazzani AR (1995) Allopregnanolone concentration in hippocampus of prepubertal rats and female rats throughout estrous cycle. J Endocrinol Invest. 18(11):853–6.

Pantazopoulos H, Gisabella B, Rexrode L, Benefield D, Yildiz E, Seltzer P, Valeri J, Chelini G, Reich A, Ardelt M, Berretta S (2020) Circadian rhythms of perineuronal net composition. eNeuro. 7(4):ENEURO.0034-19.2020.

Patel J, Fujisawa S, Berényi A, Royer S, Buzsáki G (2012) Traveling theta waves along the entire septotemporal axis of the hippocampus. Neuron 75(3):410–7.

Pineles SL, Nillni YI, Pinna G, Irvine J, Webb A, Arditte Hall KA, Hauger R, Miller MW, Resick PA, Orr SP, Rasmusson AM (2018) PTSD in women is associated with a block in conversion of progesterone to the GABAergic neurosteroids allopregnanolone and pregnanolone measured in plasma. Psychoneuroendocrinology.93:133–141.

Plappert CF, Rodenbücher AM, Pilz PK (2005) Effects of sex and estrous cycle on modulation of the acoustic startle response in mice. Physiol Behav. 84:585–94.

Reardon LE, Leen-Feldner EW, Hayward C (2009) A critical review of the empirical literature on the relation between anxiety and puberty. Clin Psychol Rev. 29(1):1–23.

Reynolds TA, Makhanova A, Marcinkowska UM, Jasienska G, McNulty JK, Eckel LA, Nikonova L, Maner JK. (2018) Progesterone and women’s anxiety across the menstrual cycle. Horm Behav. 102:34–40.

Rhodes ME, Frye CA. (2001) Inhibiting progesterone metabolism in the hippocampus of rats in behavioral estrus decreases anxiolytic behaviors and enhances exploratory and antinociceptive behaviors. Cogn Affect Behav Neurosci. 1:287–96.

Rocca WA, Grossardt BR, Geda YE, Gostout BS, Bower JH, Maraganore DM, de Andrade M, Melton LJ 3^rd^ (2018) Long-term risk of depressive and anxiety symptoms after early bilateral oophorectomy. Menopause. 25(11):1275–1285.

Romeo RD, Lee SJ, McEwen BS (2004) Differential stress reactivity in intact and ovariectomized prepubertal and adult female rats. Neuroendocrinology. 80(6):387–93.

Ross LE, McLean LM (2006) Anxiety disorders during pregnancy and the postpartum period: A systematic review. J Clin Psychiatry. 67:1285–98

Sainsbury RS, Heynen A, Montoya CP (1987) Behavioral correlates of hippocampal type 2 theta in the rat. Physiol Behav. 39(4):513–9.

Schrader AJ, Taylor RM, Lowery-Gionta EG, Moore NLT (2018) Repeated elevated plus maze trials as a measure for tracking within-subjects behavioral performance in rats (Rattus norvegicus). PLoS One. 13(11):e0207804.

Shanmugan S, Sammel MD, Loughead J, Ruparel K, Gur RC, Brown TE, Faust J, Domchek S, Epperson CN (2020) Executive function after risk-reducing salpingo-oophorectomy in BRCA1 and BRCA2 mutation carriers: does current mood and early life adversity matter? Menopause. 27:746–755.

Smith SS, Shen H, Gong QH, Zhou X (2007) Neurosteroid regulation of GABA(A) receptors: Focus on the alpha4 and delta subunits. Pharmacol Ther. 116:58–76.

Somers JM, Goldner EM, Waraich P, Hsu L (2006) Prevalence and incidence studies of anxiety disorders: a systematic review of the literature. Can J Psychiatry. 51:100–13.

Sorg BA, Berretta S, Blacktop JM, Fawcett JW, Kitagawa H, Kwok JCF et al. (2016) Casting a wide net: role of perineuronal nets in neural plasticity. J Neurosci 36:11459–11468.

Ström JO, Theodorsson A, Ingberg E, Isaksson IM, Theodorsson E (2012) Ovariectomy and 17β-estradiol replacement in rats and mice: a visual demonstration. J Vis Exp. 64:e4013.

Thorpe JB, Gould KE, Borman ED, deCatanzaro D (2014) Circulating and urinary adrenal corticosterone, progesterone, and estradiol in response to acute stress in female mice (Mus musculus). Horm Metab Res. 46:211–8.

Trombello JM, Pizzagalli DA, Weissman MM, Grannemann BD, Cooper CM, Greer TL, Malchow AL, Jha MK, Carmody TJ, Kurian BT, Webb CA, Dillon DG, McGrath PJ, Bruder G, Fava M, Parsey RV, McInnis MG, Adams P, Trivedi MH (2018) Characterizing anxiety subtypes and the relationship to behavioral phenotyping in major depression: Results from the EMBARC study. J Psychiatr Res. 102:207–215.

Vythilingum B (2008) Anxiety disorders in pregnancy. Curr Psychiatry Rep. 10:331–5.

Wakatsuki Y, Inoue T, Hashimoto N, Fujimura Y, Masuya J, Ichiki M, Tanabe H, Kusumi I (2020) Influence of childhood maltreatment, adulthood stressful life events, and affective temperaments on premenstrual mental symptoms of nonclinical adult volunteers. Neuropsychiatr Dis Treat. 16:1–10.

Wingert JC, Sorg BA (2021) Impact of perineuronal nets on electrophysiology of parvalbumin interneurons, principal neurons, and brain oscillations: a review. Front Synaptic Neurosci. 13:673210.

Wu X, Gangisetty O, Carver CM, Reddy DS (2013) Estrous cycle regulation of extrasynaptic δ-containing GABA(A) receptor-mediated tonic inhibition and limbic epileptogenesis. J Pharmacol Exp Ther. 346:146–60.

Yang S, Sylvain G, Molinaro A, Naito-Matsui Y, Hilton S, Foscarin S et al. (2020) Restoring the pattern of proteoglycan sulphation in perineuronal nets corrects age-related memory loss. bioRxiv 20.01.03.894188.

Yehuda R, Bierer LM, Andrew R, Schmeidler J, Seckl JR (2009) Enduring effects of severe developmental adversity, including nutritional deprivation, on cortisol metabolism in aging Holocaust survivors. Journal of psychiatric research, 43(9): 877–883.

Yeung M, Treit D, Dickson CT (2012) A critical test of the hippocampal theta model of anxiolytic drug action. Neuropharmacology. 62(1):155–60.

Yohn CN, Shifman S, Garino A, Diethorn E, Bokka L, Ashamalla SA, Samuels BA (2020) Fluoxetine effects on behavior and adult hippocampal neurogenesis in female C57BL/6J mice across the estrous cycle. Psychopharmacology. 237:1281–1290.

Young CK, McNaughton N (2009) Coupling of theta oscillations between anterior and posterior midline cortex and with the hippocampus in freely behaving rats. Cereb Cortex. 19(1):24–40.

Zhu J, Lowen SB, Anderson CM, Ohashi K, Khan A, Teicher MH (2019) Association of prepubertal and postpubertal exposure to childhood maltreatment with adult amygdala function. JAMA Psychiatry. 76(8):843–853.

